# Comparing Multislice Projections of MD Simulations with CryoEM Exposes Structural Prediction Errors

**DOI:** 10.64898/2025.12.09.693260

**Authors:** Arshad Mohammed, James Lincoff, Andrew Natale, Colin Ophus, Michael Grabe, Adam Frost, Frank R. Moss

## Abstract

Cryo-electron microscopy (cryoEM) is a powerful tool for atomic- and molecular-resolution structure determination, while molecular dynamics (MD) simulations are similarly powerful tools for predicting molecular trajectories. Given the challenges in estimating biomolecule dynamics with cryoEM alone, MD simulations are employed to forecast molecular motions and to interpret cryoEM reconstructions. Few methods, however, can evaluate MD predictions directly. Here, we use multislice wave propagation to project sampled snapshots of MD trajectories, either coarse-grained (CG) or all-atom (AA), into simulated cryoEM 3D reconstructions. We compared simulated and experimental images of low- and high-curvature membranes to show that MD simulations qualitatively reflect the fluidity and thus the contrast of biological membranes observed by cryoEM. MD simulations also correctly predicted bilayer dimensions for single component flat bilayers observed in cryoEM images. However, Martini3 CG-MD simulations failed to predict changes in membrane thickness induced by high curvature and with heterogeneous lipid compositions. We pinpointed the misbehavior of polyunsaturated lipid tails and cholesterol in Martini3 simulations as the main error sources contributing to inaccurate bilayer thicknesses. Our comparisons also explain membrane structure discrepancies between cryoEM and small angle X-ray scattering (SAXS). Further testing of MD predictions by direct comparisons between simulated and experimental cryoEM images should lead to the development of more accurate MD force fields.

**Statement of Significance:** Molecular dynamics (MD) simulations are frequently employed to predict the dynamics of biological macromolecules and assemblies, but these predictions remain difficult to validate experimentally. Cryo-electron microscopy (cryoEM) can be used to directly image the conformational ensemble of macromolecules, but images of Coulombic potential cannot be easily compared to snapshots of atoms from MD simulations. Here, we show that a physics-based multislice image projection algorithm accurately converts MD trajectories to simulated cryoEM 2D images and 3D reconstructions. Using this approach, we identify consistencies and discrepancies between MD simulations and cryoEM experiments. Notably, coarse-grained MD performs poorly compared to all-atom MD when simulating membranes composed of mixtures of lipids that include cholesterol and polyunsaturated lipids, providing observables for MD force field improvement.

## Introduction

Molecular dynamics (MD) simulations play a central role in investigating the structural dynamics of biological systems in realistic solution conditions (1). These dynamics are critical to the function and regulation of biomolecules but are experimentally difficult to measure. Structural biology methods such as cryo-electron microscopy (cryoEM) and X-ray crystallography provide density maps that represent the solution ensemble (cryoEM) or crystalline (X-ray) average of the measured molecules, respectively. In some cases, we can extract information about discrete conformational states with fractional occupancy from these data (2–7). Other techniques, including nuclear magnetic resonance (NMR) spectroscopy, electron paramagnetic resonance (EPR) spectroscopy, and atomic force microscopy (AFM), also provide dynamical information, but have limiting sample requirements (8–11). Molecular structures from cryoEM and X-ray crystallography have played a pivotal role in obtaining atomistic details of molecular assemblies. However, they provide limited resolutions of more fluid, disordered, and otherwise conformationally dynamic systems, including membranes, intrinsically disordered nucleic acids and proteins, colloidal condensates, and other important states of active matter.

Given the challenges of extracting dynamical information from static images, we often turn to MD simulations to reveal the motions of biological systems in the nanosecond to millisecond regime (1, 12). However, it remains challenging to falsify MD predictions (13). For example, it is not currently possible to directly compare MD snapshots of a lipid bilayer with the low-resolution bands of leaflet intensity in cryoEM images of membranes. A tool to convert MD trajectories into simulated cryoEM data would allow researchers to falsify MD simulations, provide additional observables by which MD force fields can be parameterized, and improve other applications that compare experimental and simulated data, such as ensemble- or “flexible-fit” model building into potential maps (14, 15).

Many groups have utilized approaches to convert atomic coordinates from MD into simulated transmission electron microscopy (TEM) two-dimensional images or volumes, with varying degrees of success (16–23). The simplest approach is to replace each atom with a voxel representation of its Coulomb potential, convolved with a Gaussian blur to approximate a specified resolution, and these voxel arrays can be projected into 2D images. This method is implemented in many structural biology software packages (17–19) but does not create images or volumes that recapitulate experimental data because they often neglect temperature factors, the transfer functions of the microscope and detector, the contribution of amplitude contrast, Ewald sphere symmetry breaking, or the contributions of solvent and solute molecules.

More complex approaches incorporate a multislice algorithm to model the interaction of the electron beam with all of the subject, solvent, and solute atoms, and can incorporate microscope aberrations, but this work has so far been limited to 2D image evaluation (16, 24–28). Finally, for biological lipid bilayers and peptoid membranes, groups have used MD trajectories of flat bilayers to predict the bilayer profiles of vesicles in cryoEM images (29–32). This approach has so far been limited to protein-free systems and the analysis of 2D images, rather than 3D volumes. To our knowledge, no one has reported studies of the dynamics of biological assemblies by forward mapping of MD trajectories to simulated 3D cryoEM reconstructions that can be compared directly with experimental reconstructions in 3D.

Here, we report a method of converting coarse-grained (CG, Martini) and all-atom (AA, CHARMM36) MD trajectories into simulated cryoEM 3D reconstructions. The CHARMM36 force field, coupled with a dynamics engine (GROMACS here), provides the dynamic evolution of a system by calculating the pairwise bonded and nonbonded interactions between atoms as a function of time and updating these coordinates over short time steps. While AA force fields like CHARMM36 often reproduce the dynamical behavior of molecules, they treat atoms classically and employ a variety of approximations to make computation feasible. The CG Martini force field is similar except that groups of approximately four heavy atoms are replaced by a single bead with empirically determined potentials modeling the interactions between beads. CG force fields such as Martini3 offer tremendous enhancements in sampling speed over conventional AA models like CHARMM36. They do so by smoothing the energy landscape to allow much longer time steps and reducing the computational burden by decreasing the particle number, but these computational gains necessarily trade-off chemical fidelity.

In this work, we consider three model systems: flat lipid bilayers, curved lipid bicelles, and a nanotube of membrane induced by a copolymer of human ESCRT-III proteins, CHMP1B and IST1. In our previously reported cryoEM maps of the copolymer assembly, the proteins are resolved to near-atomic resolution, while the lipids in the membrane nanotube are smeared into continuous density for each leaflet due to the incoherent averaging of fluid molecules in the 2D plane of the leaflet (33, 34). Here, we sought to predict and measure microsecond membrane dynamics and nanosecond lipid-lipid and lipid-protein interactions with CG- and AA-MD and to compare these with experimental cryoEM observables.

To directly compare MD simulation trajectories with experimental cryoEM reconstructions, we developed a pipeline that uses an adaptation of the multislice method (27) to generate projected TEM images from MD trajectory snapshots, followed by back-projection 3D reconstructions using RELION (35). We generated simulated 3D reconstructions that include ns-scale AA and μs-scale CG dynamics, microscope aberrations, as well as solvent and solute contributions. Encouragingly, the computed Coulombic potential maps qualitatively and quantitatively match many aspects of the experimental data. However, the simulations underestimate membrane thickness in some instances. This discrepancy is particularly strong in the highly curved, four-component membrane nanotubes bound by ESCRT-III proteins.

By simulating zero-curvature bilayers with atomistic and CG-MD, we show that discrepancies result from cholesterol and polyunsaturated lipids, whose localizations within the bilayer differ strongly between simulations using the two force fields. With Martini force fields, polyunsaturated tails have a higher likelihood of bending towards the bilayer surface, while cholesterol is more likely to localize to the midplane of bilayers. These results suggest that the Martini force field parameterization does not function well for certain lipids and may need to be reparametrized based on observed properties in bilayers with diverse compositions and geometries. Lastly, we use the methodology described here to explain the previously observed differences in apparent bilayer thicknesses measured by cryoEM and SAXS. We show that these differences result from the anisotropic shape of the Coulombic potential profile of the bilayer and the disparate resolutions of the two techniques.

## Materials and Methods

### Materials

Lipids were from Avanti Polar Lipids (Alabaster, AL, USA). TEM grids were from Quantifoil (Großlöbichau, Germany). All other reagents were from ThermoFisher Scientific (Waltham, MA, USA) and were used as supplied.

### Experimental data

Experimental cryoEM reconstructions of CHMP1B/IST1 were accessed via the EMDB from EMD-28700. Solution SAXS data were from Ref. (36).

### Coarse-grained molecular dynamics simulations of tubules

Four sets of structures of IST1-CHMP1B copolymers around a membrane tubule were generated in CG-MD simulations for back-mapping to AA representation. We refer to the structures as tubule Sets 1-4, summarized in Table 1. Data Set 1 was sourced directly from our previous work on the system (33)—10 evenly spaced timepoints from one replicate of a Martini 2.2 simulation comprising two turns of the CHMP1B-IST1 protein coat (referred to as Simulation Set 1 in that work). Along with this two-turn dataset, we generated three additional sets of four-turn structures (Sets 2-4).

**Table 1.**
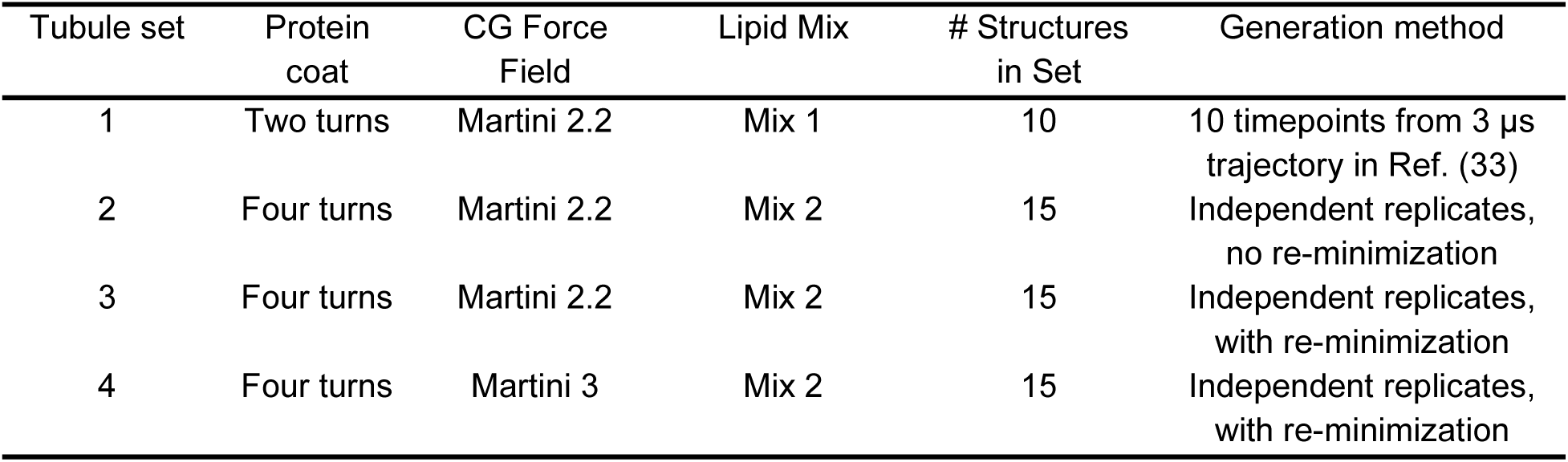
Summary of ESCRT-III tubule CG simulation datasets.

Sets 2-4 each consisted of fifteen replicates. The structure setup, initialization, and run procedure largely followed our previous simulations on the same system (33) with reduced MD runtimes since our aim was to generate structures for conversion to AA rather than study long-time CG dynamics. System setup was done using GROMACS and Martini functions and scripts, and MD was run using GROMACS 2020.6 (37).

Initial structures were built by converting the cryoEM structure of the IST1-CHMP1B copolymer to Martini 2.2 (38) or Martini3 (39) CG representation using the Martini *martinize* and *martinize2* scripts (40). To maintain the protein structure during MD, all protein bead positions were fully restrained throughout, and no elastic network was applied, as described previously (33). The CG protein was centered in a 34 x 34 x 34 nm cubic box, aligned such that the membrane tubule would form through the box center along the z axis. Lipids were then added as in our previous work (33), placed within the cylindrical volume where the tubule forms during spontaneous assembly. Pre-determined leaflet compositions were generated by dividing the cylindrical volume into inner leaflet and outer leaflet regions and adding specified numbers of each lipid type to each region to match the cryoEM-determined membrane leaflet compositions (Table 2, comprises Mix 2 from SI Table 1). Slightly more lipids were added in the Martini3 simulations than for Martini 2.2, following tuning of the lipid packing for the new force field. Structures were then solvated with Martini water, ionized to 150 mM NaCl, and energy minimized with 2500 steps of a steepest descent algorithm.

**Table 2.** Lipid counts per leaflet for four-turn simulated structure initialization.

|  | Lipid count |  |  |  |
| --- | --- | --- | --- | --- |
|  | Martini 2.2 |  | Martini3 |  |
|  | Inner leaflet | Outer leaflet | Inner leaflet | Outer leaflet |
| SDPC | 185 | 336 | 187 | 339 |
| POPS | 351 | 503 | 355 | 508 |
| CHOL | 273 | 305 | 276 | 308 |
| PIP<sub>2</sub> | 166 | 381 | 168 | 385 |

The above coordinate generation was repeated five times for each simulation set, to produce five distinct initial configurations of randomized lipid positions. Each configuration was then run through an equilibration MD step three times with randomized velocity initialization to produce the fifteen distinct structures per dataset. This equilibration is an abbreviated version based on our previous work (33), as there is no need to perform long-time equilibration of leaflet compositions. Main simulation parameters conform to standards for simulating with Martini and are maintained in all MD stages: temperature of 320 K maintained with the velocity rescaling thermostat, timestep of 25 fs, reaction-field electrostatics, and a 1.1 nm Coulomb cutoff.

Equilibration was run in three steps. First, there is a 500 ns segment for spontaneous assembly of the lipid tubule, during which a 2 nm radius pore was maintained with a cylindrical, flat-bottomed repulsive restraint potential applied to the lipid tails, as was done previously (33). The pore allows water to freely enter and exit the tubule lumen to reach proper internal solvation. The second stage is a brief 25 ns intermediate step where the pore radius was reduced by half, before the final 100 ns step where the pore is fully closed. This total equilibration is short to limit lipid flip-flop at the pore, such that target leaflet compositions were maintained. The first equilibration step used a semi-isotropic Berendsen barostat, due to its greater stability, before switching to semi-isotropic Parrinello-Rahman for the last two steps. Throughout equilibration, a reduced force constant of 20 kJ mol^-1^ nm^-2^ was applied for the protein position restraints, versus the default value of 1000 used previously for the two-turn structures, as this greatly improves simulation stability given the nearly 27,000 protein residues to restrain. Following MD, the final frame for each replicate was energy minimized again, with 10,000 steps of steepest descent and a 1000 kJ mol^-1^ nm^-2^ force constant on the protein, to return the protein bead coordinates to the exact initial configuration following conversion from AA (Sets 3 and 4). One set of structures was additionally back-mapped before this re-minimization (Set 2) to observe the effect of reduced protein alignment on the multislice pipeline.

### All-atom back-mapping

Standard Martini *backward* protocols were followed to generate AA CHARMM36-parameterized configurations from energy-minimized CG simulation snapshots (41). The standard protocol includes three basic stages to produce an atomic configuration: first, mapping atoms onto CG bead positions; second, energy minimization with only bonded potentials to get correct molecular geometries; and third, energy minimization with non-bonded potentials turned on to obtain correct overall molecular packing. Following the bonded-only minimization, we applied an in-house Python script that searches for and relaxes atomic clashes of < 0.5 Å within the set of ∼730,000 protein and lipid atoms. This removes clashes that would otherwise cause the non-bonded minimization to fail, typically ∼500 clashes per structure.

### All-atom tubule simulation

A final set of ESCRT-bound tubule structures was generated from an all-atom trajectory that was initialized using one of the backmapped structures from Set 4 for starting coordinates. The simulation was run with Gromacs 2020.6 using a 2 fs timestep and was run for 200 ns.

Temperature was maintained using the stochastic velocity rescale thermostat at 303.15 K, and the semi-isotropic Parinnello-Rahman barostat maintained pressure at 1 atm. Protein heavy atoms were restrained to maintain the cryo-EM resolved complex structure. Two sets of simulation frames, spanning the 125-150 ns and 175-200 ns segments and sampled per nanosecond, were evaluated.

### Multislice algorithm

The cryoEM simulations were performed using the multislice method, originally proposed by Cowley and Moodie (42). We followed the methods outlined by Kirkland, implemented in MATLAB (26). First, we import the atomic coordinates from the PDB file format. We rotated, tiled, and cropped the resulting cells to create the desired simulation geometry. Next, we computed the projected potentials using a lookup table method (43) and used the multislice method to propagate plane waves through these projected potentials to produce exit waves. Finally, we applied the microscope transfer function to these exit waves, including defocus and spherical aberration to match the experimental conditions. We assumed a 1 Å information limit, applied using a temporal coherence envelope (44). and calculated noisy output images using Poisson statistics with the appropriate electron dose. For the ESCRT-III filament simulations, we used a final pixel size of 0.835 Å, an electron dose of 65 e^-^/A^2^, and accelerating voltage of 300 kV to match the cryoEM experimental parameters. For the protein-free lipid simulations, we used a final pixel size of 1 Å, an electron dose of 65 e^-^/A^2^, and accelerating voltage of 200 kV, also to match experimental conditions. The efficiency of the multislice algorithm scales inversely with the number of atoms, so we attempted to optimize the set of atoms used to generate projections. We explored using explicit solvent versus Poisson noise and found no advantage to the explicit solvent, so we utilized the Poisson noise model for computational efficiency (Fig. S1A). We also investigated whether excluding hydrogen atoms affected the projections. Exclusion of hydrogen atoms resulted in a small (∼ 1 Å) apparent thickening of a POPC bilayer (Fig. S1B). For computational efficiency, we chose to exclude hydrogens from multislice image projection.

### Processing simulated data

Simulated ESCRT filament projection images from the multislice program were processed as described previously for experimental data following preprocessing as follows. RELION requires square particle images with a mean of 0 and a standard deviation of 1 in an MRCS format (45). *Tif2mrc* in IMOD (46) was used to concatenate simulated particles into an MRCS stack. The relion_preprocess tool was then used to adjust the mean and standard deviation of the particle pixel values per RELION conventions. Finally, a Python script was used to write a STAR file with the appropriate priors for filamentous particles.

After preprocessing, the particle stacks were processed as previously described (33) in RELION3.1.4 with the following parameters. 2D classification was performed with one class, a tube diameter of 250 Å, and a τ value of 2. A 3D_autorefine was subsequently performed with the aligned particles with a featureless cylinder as a reference, 25 Å inner filament diameter, 260 Å outer filament diameter, 17 asymmetric units per turn, 20.8 ° initial twist, 3.18 Å initial rise, and central 30% of the helix. For both 2D classification and 3D_autorefine, the first peak of the CTF was ignored. Power spectra for 2D averages were calculated in RELION. The handedness of the simulated 3D reconstructions was inverted using the relion_image_toolbox tool when necessary.

Simulated flat bilayer projection images were analyzed in the same manner except they were only subjected to 2D classification. Prior to processing in RELION, the particle stacks were low-pass filtered to various resolutions with the relion_image_toolbox tool. In 2D classification, a τ value of 0.1 was used.

### Data visualization

The final simulated reconstruction of ESCRT filaments was visualized by summing the central 10 horizontal and vertical slices. For radial profiles only, the volumes were low-pass filtered to 6 Å. Radial profiles were created by summing the central 30% of the slices and computing the radial profile with the Radial Profile Angle plugin in FIJI (47). The full z and y projections of the volumes were also generated for visualization of both experimental and simulated volumes.

### Molecular dynamics simulations of flat lipid bilayers and curved bicelles

Both CG and AA simulation systems of planar lipid bilayers were prepared for pure POPC, a quaternary mixture with SDPC (58% SDPC, 18% POPS, 18% cholesterol, 6% PIP_2_), a quaternary mixture with POPC (58% POPC, 18% POPS, 18% cholesterol, 6% PIP_2_), and a cholesterol-free ternary mixture with SDPC (71% SDPC, 21% POPS, 8% PIP_2_) (see Table S1 for lipid mixtures). An additional AA system was prepared with pure DPhPC lipids. The PIP_2_ species used for all simulations was dioleoyl-PI(3,5)P_2_. To set up the simulations, initially 800 lipids were randomly inserted into a box of dimension 16×16×13 nm using the GROMACS ‘insert-molecules’ tool. The remaining volume of each box was filled with water molecules, and then 150 mM Cl^-^ and neutralizing Na^+^ ions were added by randomly replacing water molecules. For AA simulations, the CHARMM36 force field was used for lipids (48), the TIP3P water model was used with compatible ion parameters (49, 50), and two atomic mass units (amu) of mass were repartitioned to each lipid hydrogen atom from its bound heavy atom to enable simulations with a 4 fs timestep (51). CG simulations used the Martini3 force field for lipids, water, and ions (39). We prepared and ran two variants of each Martini3 bilayer/bicelle simulation; the first set used parameters and lipid topologies available upon the original release of the Martini3 force-field, obtained from https://cgmartini.nl/ (referred to as Martini3 in the text), while the second set used updated parameters and lipid topologies resulting from recent lipid-tail reparameterization efforts (referred to as Martini3.2 in the text) (52). All lipid species used the same number of coarse-grained beads in both sets of simulations; however, in some cases bead sizes and types change in the reparametrized lipids.

Planar bilayer simulations were run with GROMACS 2022.5/6 (37). Each system was minimized for a maximum of 5000 steps using a steepest descent algorithm, and then systems were equilibrated in the NPT ensemble for 50 ps with no restraints. Following this equilibration, lipids were driven into a planar bilayer over the course of 2 ns via application of a 50 kJ mol^-1^ nm^-2^ potential acting in the z-dimension with a minimum at the center plane of the simulation cell in z, applied to the terminal carbon atom or CG bead of one of the fatty acid tails of each lipid, or C27 for cholesterol. After driving bilayer formation for 2 ns, each system was equilibrated for 500 ns (AA) or 1500 ns (CG) without restraints at 310 K and 1 atm pressure with a 4 fs (AA) or 10 fs (CG) timestep using a leap-frog integrator, a stochastic velocity rescaling thermostat (‘tcoupl=v-rescale’, AA:‘tau-t=1.0’, CG:’tau-t=2.0’), and a stochastic cell rescaling barostat (‘pcoupl=c-rescale’, ‘pcoupltype=semiisotropic’, AA:‘tau-p=5.0’, CG:‘tau-p=12.0’).

For CHARMM36 AA simulations, short range nonbonded interactions were cut off at 1.2 nm, with force-switching applied to smoothly turn off Lennard-Jones interactions in the range 1.0-1.2 nm (53), long range electrostatic interactions were calculated with the particle mesh Ewald method, and non-water bonds to hydrogen atoms were constrained with the LINCS algorithm. For Martini3 CG simulations, short range nonbonded interactions were cut off at 1.1 nm, long range electrostatic interactions were treated with the reaction field method, and force field specified bonds were constrained with the LINCS algorithm. After the equilibration phase, each trajectory was run for an additional 1500 ns (AA) or 2000 ns (CG) production phase, from which samples were drawn for analysis of membrane structural properties.

Simulations of bicelles with a 3.7 nm radius of curvature were run with GROMACS 2022.5 compiled with support for PLUMED 2.8.3 (54) to enable the use of the EnCurv method for constraining membrane curvature (55). Martini3 simulation systems of curved bicelles were derived from snapshots of the corresponding planar bilayer simulations. In each case, the initial lipid configuration was extracted from the source trajectory, placed in a new box that was larger in the x and z dimensions to accommodate the bicelle and then re-solvated with GROMACS tools. Each system was equilibrated from zero curvature to a final radius of curvature of 3.7 nm stepwise over 20 ns and then equilibrated for an additional 2000 ns. After this, each simulation was run for another 2000 ns production phase from which samples were drawn for analysis.

Three AA replicate curved bicelle simulations were constructed for each lipid mixture from back-mapped CG configurations spaced 250 ns apart in the curved bicelle CG trajectories. Each trajectory was equilibrated for 50 ns, followed by a 105 ns production run from which samples were drawn for analysis (total 315 ns production per lipid mixture). When applying EnCurv, 150 radial bins were used, and biasing forces were applied to all lipid CG particles or AA heavy atoms. Other simulation run settings were the same as planar bilayer trajectories, with the exception that pressure was controlled with an anisotropic Parrinello-Rahman barostat, allowing only changes in the y-dimension of the simulation cell, with x and z fixed.

CG configurations of planar and curved membranes were back-mapped to AA configurations using a custom implementation of the ‘backward’ procedure (56), followed by re-solvation of systems with GROMACS tools, minimization, and a short 85 ps re-equilibration simulation. During re-equilibration, harmonic positional restraints of 50 kJ mol^-1^ nm^-2^ were applied to a subset of atoms in each all-atom lipid based on the positions of the CG beads from which they were derived to relax the aqueous phase while maintaining the lipid/membrane conformations from the CG structure in the AA back-mapped structures.

### Experimental cryoEM images of lipid vesicles

CryoEM images of lipid vesicles were processed as described previously (36). Briefly, small unilamellar vesicles (SUVs) were prepared by the extrusion method. 300 nmol of the desired lipid or a lipid mixture were added to a glass scintillation vial from lipid stocks dissolved in chloroform. The solvent was removed with a gentle stream of nitrogen while rotating the vial. The lipid film was placed in a vacuum desiccator under house vacuum for 2 hours. The lipid film was resuspended by adding 250 μL of TBS and vortexing. The resulting multilamellar suspension was extruded 31 times through a polycarbonate membrane with 50 nm pores using an Avanti mini extruder. 4 μL of the SUV dispersion was applied to a glow discharged Quantifoil holey carbon grid, blotted with 0 force for 4 seconds at 19 °C under 100% humidity, and plunge-frozen in liquid ethane using a Vitrobot Mark IV.

The grids were imaged with a Glacios 200 keV microscope equipped with a Gatan K2 detector. The raw movies were processed with RELION3.1.4 (35). The movies were motion corrected with MotionCor2 in RELION (57). Overlapping regions of the bilayers were picked with the manual picking tool, extracted, CTF-corrected, and 2D aligned and averaged. The 2D classes with good alignment were reclassified into a single 2D class with τ set to 0.1. The electron scattering profiles of the flat bilayers were calculated by taking a line profile with a width of 10 px of the 2D classes in FIJI. The electron scattering profiles of the curved bicelles were calculated with the Radial Profile Angle plugin in FIJI using an angle of 90 ° centered on the bicelle.

## Results

### Simulation of ESCRT-III-bound membrane nanotubes

To develop a methodology to quantitatively compare MD simulations to experimental cryoEM data, we started by conducting CG-MD simulations with a system consisting of ESCRT-III proteins, CHMP1B and IST1, assembled in a copolymeric filament around a lipid bilayer nanotube. We chose this system as it contains both a relatively rigid protein assembly and a dynamic lipid bilayer assembly. The dynamic bilayer requires explicit treatment of lipid motion to accurately compare simulation with experiment. We assembled and simulated the system in a manner similar to our previous report (33) except that we increased the simulation size to include four turns of the helix to better match the size of the experimentally reconstructed volume. The protein coat is restrained in these simulations to maintain the cryoEM-resolved structure. We generated 15 structures from independent simulations using Martini 2.2 or Martini3, and then back-mapped the final frame from each CG-MD simulation to an AA representation using the backward Python script (56). We reasoned that using independently generated structures would ensure that the bilayer conformations were distinct, effectively mimicking the averaging across filament segments in experiments that produces smooth densities for the tubule.

We next sought to convert each series of 15 independent MD snapshots to a series of physically realistic simulated cryoEM projection images. We chose a multislice algorithm modified from the MATLAB version of the Prismatic program due to its computational efficiency, physics-based simulation approach, and ability to incorporate multiple scattering and microscope aberrations (27). In addition to sampling independent MD simulations, we also sampled multiple views of the filament by rotating the atomic coordinates around the filament axis in 1 ° increments. This procedure resulted in 5400 simulated cryoEM projection images, sampling μs of dynamics and the full range of rotational views. We created projection images with -1 – -2 μm defocus, 300 keV, and 65 electrons/Å^2^ to match the microscopy parameters. RELION cryoEM processing software requires square particle boxes, a mean intensity of 0, and an intensity standard deviation of 1, so we next preprocessed the simulated particles to meet these requirements. Finally, we created a star file, imported the simulated particle stack into RELION, and processed the stack as we processed the experimental data (see Fig. 1 for the workflow).

**Figure 1.**
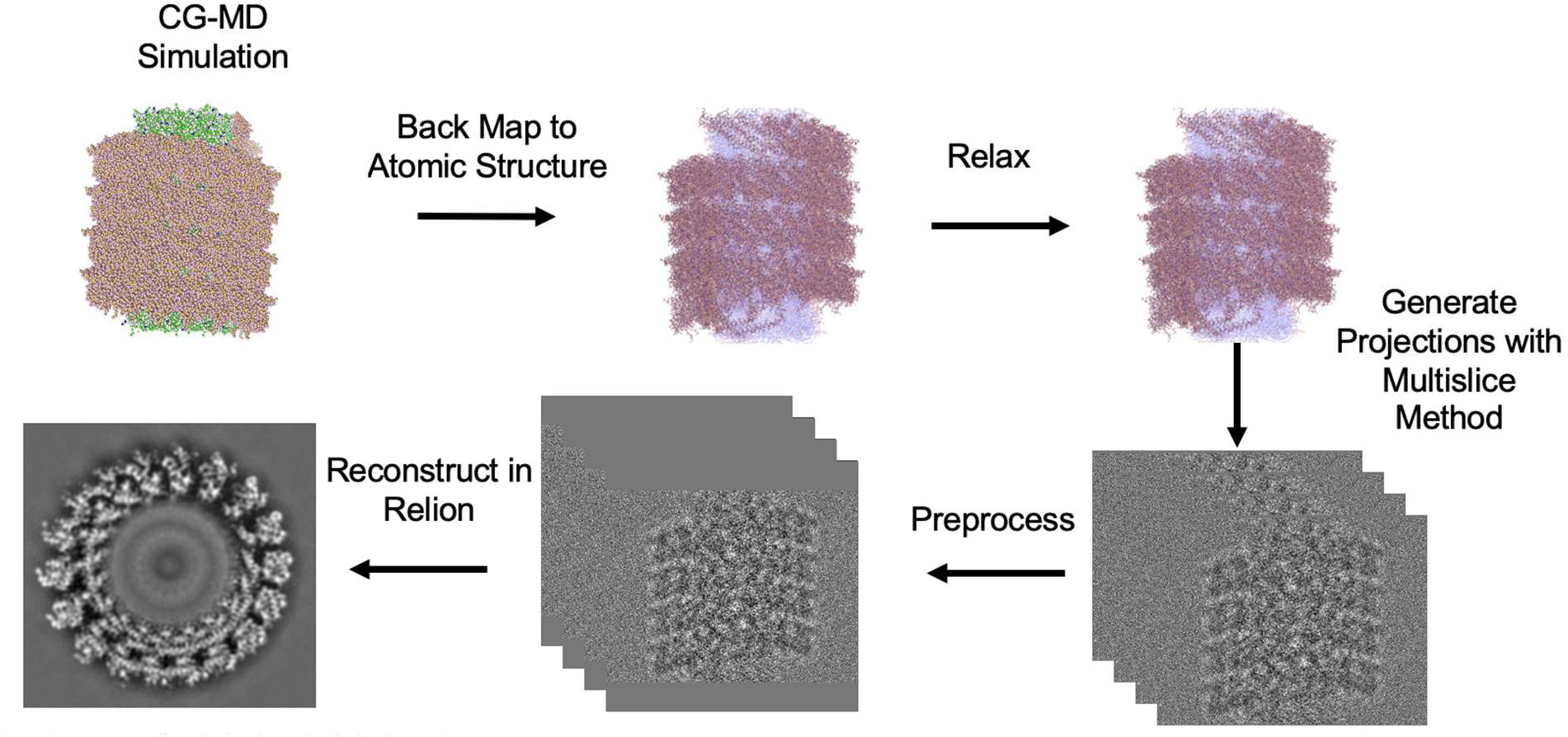
Workflow for converting CG-MD simulations to simulated cryoEM volumes. CG-MD simulation frames were back-mapped to atomistic structures and relaxed to resolve atomic clashes. Next, Coulombic potential maps were generated with the multislice wave propagation method. The output images were padded by adding pixels with the mean solvent intensity to obtain square images, cropped and binned by 2, and concatenated to form a stack. Stacks were imported to Relion3.1.4 for 2D classification and 3D reconstruction to generate a density map that includes the effects of processing experimental data.

We first qualitatively compared the simulated projection images (Set 3) to the experimental ones (Fig. S2). The 4-turn simulated filaments are not continuous across the length of the box but are otherwise similar to the experimentally derived images. We note that the signal-to-noise ratio appears higher in the simulated images. Of the potential sources of this discrepancy, we judge that ice thickness is likely the most significant. The vitreous ice in the experimental sample is >∼100 nm thick, while the simulated solvent is the thickness of the MD simulation box, ∼33 nm. Additionally, the experimental sample contains both unpolymerized protein and denatured protein at the air-water interface that contribute noise (58).

After we confirmed that the multislice algorithm could generate a distribution of 2D projections from snapshots of MD generated atomic coordinates, we processed the stack of calculated particles with RELION3.1.4 using the same workflow used to process experimental data. Specifically, we aligned and averaged the particles in 2D, followed by 3D autorefinement in RELION. Notably, the simulated 2D averages contain Fresnel fringes around protein density that result from incomplete correction of the contrast transfer function and are a recognizable feature of cryoEM data (Fig. 2). Importantly, creating projection images from multiple MD snapshots resulted in blurred intensity for each leaflet of the fluid lipid bilayer, qualitatively matching the experimental data. This agreement indicates that the number of snapshots was sufficient for the lipids to fully decorrelate. We additionally tested a series of snapshots generated every 300 ns from a single 3 μs length CG trajectory with a two-turn protein coat (10, Set 1), achieving similar decorrelation of the lipid density (Fig. S3).

**Figure 2.**
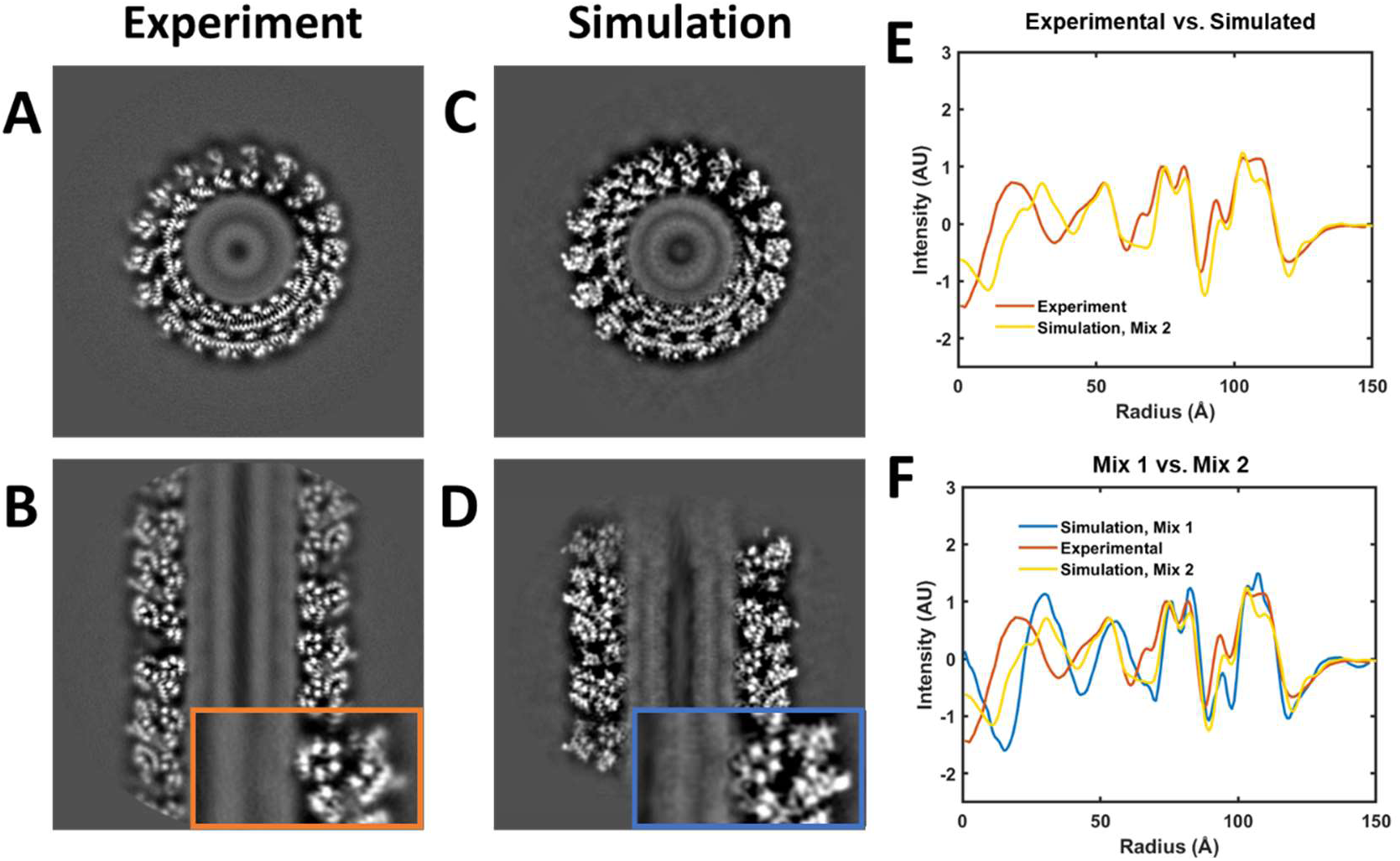
Multislice image simulations of MD snapshots generate realistic protein but not membrane reconstructions. Gray scale z projections of central 10 slices of the CHMP1B/IST1 copolymer with horizontal (A and C) and vertical views (B and D) for the experimental 3D reconstructions versus the corresponding views of the multislice simulated volume. Insets show detailed views of the CHMP1B-bilayer contact sites. E) Radial potential profiles of experimental versus CG-MD simulated native protein-coated lipid nanotubes (Mix 2). F) Radial potential profiles of experimental versus simulated volumes with different concentrations of cholesterol, SDPC, POPS, and PIP_2_ (Mixes 1 and 2).

We next examined the radial potential profiles of the membranes and protein filaments to quantitatively compare simulation with experiment. Comparing the simulated and experimental profiles in Fig. 2E, the multislice of MD snapshots workflow generates accurate intensities for each lipid bilayer leaflet relative to the protein intensity. However, the inner leaflet peak is ∼9 Å too far away from the center of the nanotube, yielding a simulated bilayer that is significantly thinner than the experimental one. Importantly, membrane thickness consistency was ambiguous when simply “fitting” coordinates from MD snapshots into the experimental cryoEM volume because it is unclear where the leaflet peaks should be relative to the time-averaged position of the distinct chemical moieties of lipids, each with a distribution of positions in the leaflet (Fig. S4). The Martini developers later released the reparametrized Martini3 to better match some experimental datasets (39), so we also simulated the ESCRT-III filament with Martini3, which resulted in a similar, but slightly thinner bilayer (Fig. S5).

Lipid composition is known to affect bilayer thickness through the length and molecular conformations of the acyl tails. In our CG-MD simulations (Fig. 2E), we used the lipid composition in the filaments that we previously estimated by measuring the signal from brominated lipids (Mix 2) (33). We next examined the sensitivity of simulated bilayer thickness to the ratio of the four lipids in the mixture by comparing the simulation with Mix 2 to a simulation with a different mixture (Mix 1). We used a previously generated CG-MD simulation with a very different ratio of the four lipids (Set 1, Mix 1) and converted this MD simulation to a simulated volume. As shown in Fig. 2F, making a large change to the lipid composition resulted in only a small increase in the bilayer thickness and a large change in the relative intensities of the leaflet peaks (Fig. 2F, Mix 1 vs. Mix 2). The small change in thickness is not surprising as it should require a completely different set of lipids to account for the 9 Å change in bilayer thickness, which is approximately the same thickness change as moving from a pure di14:1PC bilayer to a pure di22:1PC bilayer and much greater than the difference in thickness between ordered L_o_ and disordered L_d_ phases (29). In contrast, the lipids in our mixtures have acyl tails that vary between 16 and 22 carbons in length, with the majority (63%) of tails containing 18 carbons. Therefore, it is unlikely that our uncertainty in the lipid composition found in the tubules accounts for the discrepancy in bilayer thickness observed between our Martini simulations and experiment. Rather, we hypothesized that the Martini force field parameters, which were empirically determined for flat, protein-free, homogenous bilayers, fail under high membrane curvature in a heterogenous setting (59–61).

### Simulations of flat bilayers

To test this hypothesis, we turned to MD simulations of flat bilayers, which are practical to simulate with both AA and CG force fields. Additionally, the thickness of low-curvature lipid bilayers is straightforward to measure with small angle X-ray scattering (SAXS) and cryoEM. We simulated five different lipid bilayer patches of 800 lipids each composed of pure POPC, pure DPhPC, Mix 1, Mix 3, and Mix 4 using CG Martini3 and AA CHARMM36, for 2000 ns or 1500 ns, respectively (Fig. 3A). Snapshots of each simulation were sampled every 100 ns. Each of these simulation snapshots were converted to multislice simulated projection images as described for the ESCRT filaments. These simulated projections were aligned and averaged in 2D in RELION3.1. For pure POPC, pure DHPC, and Mix 1, we also prepared small unilamellar vesicles (SUVs), vitrified them, and imaged them with cryoEM. For POPC and DPhPC, we also compared the simulated and experimental cryoEM 2D averages with bilayer thickness measurements from published SAXS studies of SUVs (36, 62).

**Figure 3.**
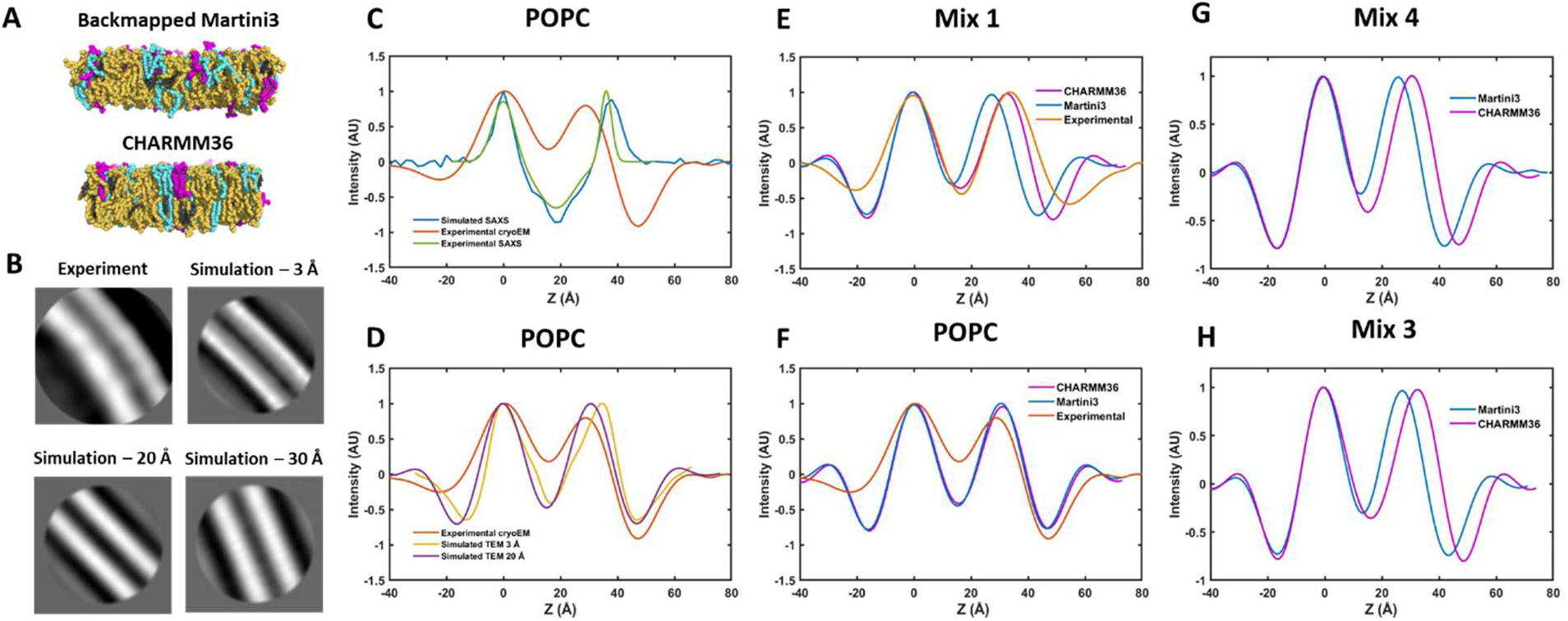
CHARMM36 and Martini3 predict flat bilayer thickness accurately, but only if cholesterol or polyunsaturated lipids are absent. A) Snapshot of MD simulations of Mix 1 bilayers. B) Multi-slice wave propagation simulated cryoEM images of snapshots of a CHARMM36 MD simulation of a POPC bilayer as in (A) compared to an experimental 2D average of cryoEM images of POPC vesicles, with specified low-pass filtering. C) Bilayer profiles of simulated (Martini3) and experimental SAXS compared to experimental cryoEM. D) Multislice simulated cryoEM bilayer profiles at different resolutions compared to an experimental cryoEM bilayer profile of POPC. E-F) MD-simulated bilayer profiles compared to experimental bilayer profiles for Mix 1 (E) and POPC (F). E-F) Comparison of MD simulated bilayer profiles for Mix 4 (G) and Mix 3 (H).

When we compared the bilayer profiles from the Martini3-simulated POPC 2D average to the experimental cryoEM POPC bilayer profile, the simulated profile was 5.6 Å thicker as measured by the peak-to-peak distance (Fig. 3B-C, Table S2). However, we also noticed that the simulated bilayer profile had higher resolution features than the smooth peaks measured by cryoEM, indicating that two 2D averages had different resolutions. The features in the simulated profile matched features seen in high-resolution SAXS studies well, e.g. a shoulder on the interiors of the phosphate peaks approximately at the location of the glycerol backbone. We therefore low-pass filtered the simulated POPC particle stacks to various resolutions before reprocessing in RELION (Fig. S6). At 20-30 Å filtering, the smooth leaflet peaks were well matched to the shape of the experimental data (Fig. 3D). Due to the asymmetric shapes of the leaflet peaks arising from the chemical structures of the lipids, this low-pass filtering resulted in anisotropic peak broadening and an apparent thinning of the bilayer (as measured by peak-to-peak distance). The simulated POPC bilayer was then only 2.6 Å thicker than the experimental data (Table S2).

We then computed the simulated SAXS profile of the POPC Martini3 simulation snapshots with the Density Profile Tool in VMD and compared the output to experimental SAXS data (63, 64). The simulated flat bilayer was 1.7 Å thicker than the experimental SAXS data, indicating that the MD simulation, or the processing pipeline, somewhat overestimates the bilayer thickness (Fig. 3C). Notably, the simulated SAXS and cryoEM profiles had somewhat different shapes (Fig. 3C vs. 4d), perhaps reflecting the different scattering factors of electrons versus X-rays or differences in sensor sensitivities. After matching the resolutions of the simulated and experimental cryoEM data, the bilayer thicknesses agreed within 1 Å. The results from CHARMM36-simulated POPC were essentially identical to Martini3-simulated POPC, indicating that for single-component, flat bilayers, the Martini3 coarse-grained model is remarkably consistent with CHARMM36 AA MD (Fig. 3F). We further compared the CHARMM36-simulated, low-pass filtered 2D average for DPhPC and Mix 1 with experimental cryoEM data (Fig. S7). When similarly low pass filtered, DPhPC (Fig. S7D) and Mix 1 (Fig. 3E) bilayer thickness values from CHARMM36 simulations agreed well with experimental cryoEM data. On the other hand, Mix 1 bilayer thickness predicted from low pass filtered Martini3 simulations produced a significantly thinner (6 Å) bilayer profile than the CHARMM36 simulations and cryoEM data (Fig. 3E), suggesting that the membrane composition plays an important role in the membrane thinning observed in the Martini tubule simulations. Simulating the bilayers without cholesterol (Mix 3) and with POPC instead of SDPC (Mix 4) reduced but did not eliminate the discrepancy in thickness between CHARMM36 and Martini3 (5 Å and 3 Å for Mix 3 and Mix4, respectively) (Fig. 3G-H, S7, and S9).

**Figure 4.**
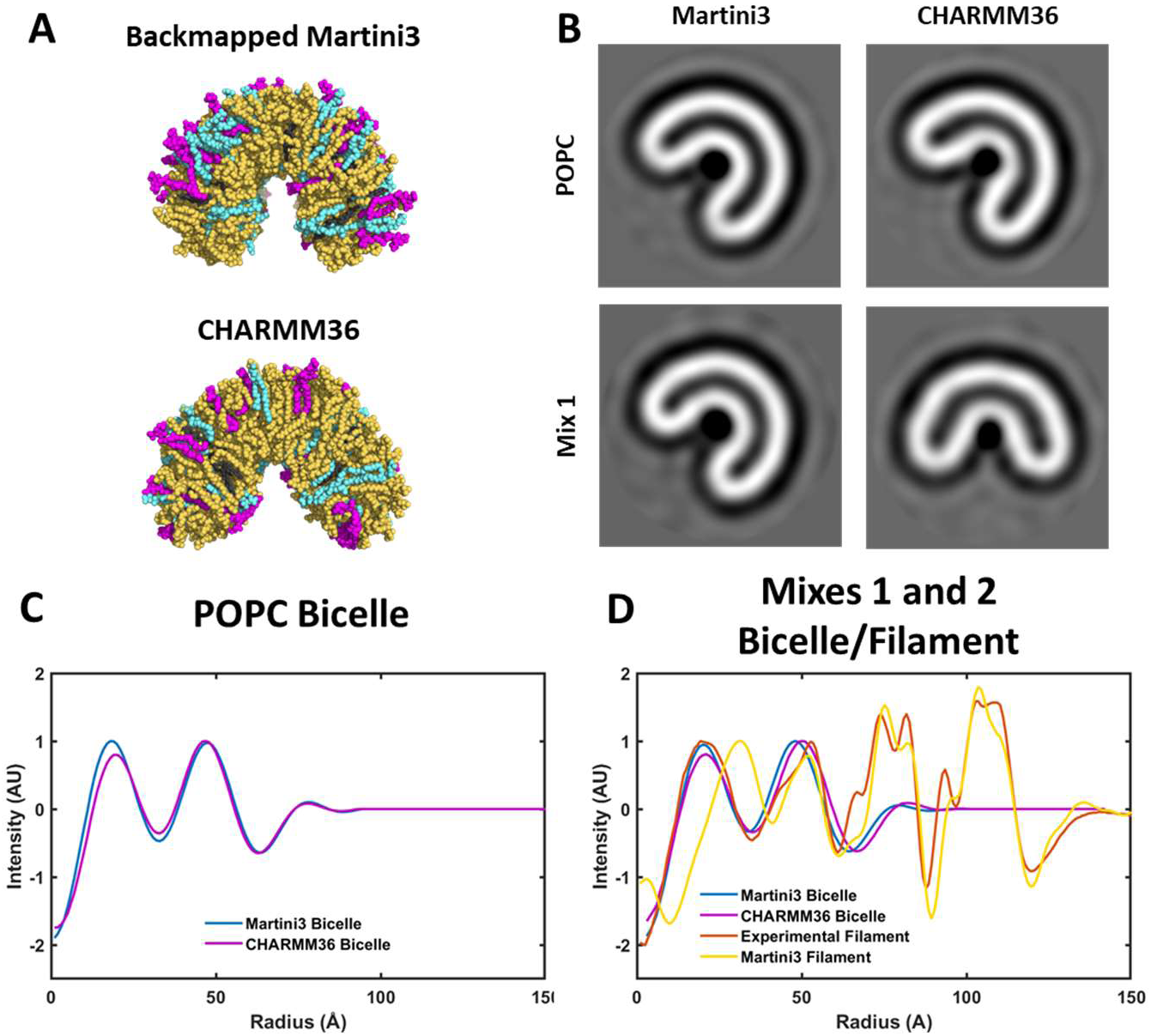
Lipid bilayers respond differently to curvature with Martini and CHARMM36. (A) Snapshots of MD simulations of curved bicelles of Mix 1. (B) Multislice wave propagation simulated cryoEM images of curved bicelles from simulations using Martini3 (left) or CHARMM36 (right) with pure POPC (top) or the quaternary lipid mixture (bottom). (C) Radial potential profiles of the central 90 ° of the half tube for the POPC system show bilayer thinning in CHARMM due to pinching. (D) Radial potential profiles for curved bicelles of Mix 1 with the experimental cryoEM and Martini3-generated data for the ESCRT filament (Mix 2). Both Martini3 and CHARMM36 (Mix 1) protein-free bicelles more closely match the experimental bilayer thickness from the copolymer filament.

### Simulation of curved bicelles

To further investigate why the Martini force field failed to match the bilayer thickness in the case of the high-curvature ESCRT-bound membrane nanotube, we used a recently reported protocol to simulate protein-free lipid bicelles with the same curvature as the ESCRT-bound tube (Fig. 4A) (55). This method allows for simulation at fixed curvature by application of a harmonic restraining potential on lipid heavy atoms, while not restricting transverse diffusion or lipid flip-flop at the bicelle edges. As in the case of flat bilayers, when the CHARMM36 (initially back-mapped from Martini3 snapshots) and Martini3 POPC bicelle simulations were converted to multislice-simulated cryoEM 2D averages, they agreed well (Fig. 4B-C) suggesting that the mechanics of high curvature are similar, at least with pure POPC bilayers.

Notably for both CG and AA MD, the curved POPC bicelle was thinner than the flat POPC bilayer, in agreement with previous work (Fig. S8) (33, 55). However, CHARMM36 and Martini3 predicted strikingly different shapes when we simulated curved bicelles of Mix 1. Martini3 predicted a smoothly curved half tube, while CHARMM36 predicted a “pinched” shape with a flatter region in the middle of the half tube. In this flatter region, the CHARMM36 bilayer was thinner than at the edges (Figs. 4B, S7, and S8C).

From the different predicted bicelle shapes, it was clear that the membrane mechanics of the mixed membranes are quite different in the two force fields, with CHARMM36 predicting a kink in the bicelle. Martini3 predicted a bilayer thickness (as measured by the peak-to-peak distance in the bicelle 2D average) that was more similar to the experimentally measured thickness in the ESCRT filament than in the Martini3 ESCRT filament simulation, suggesting that inaccuracies in protein-lipid interactions may contribute to the excess bilayer thinning in the Martini3 ESCRT filament simulation (Fig. 4D). Martini has been reported to overestimate the strength of membrane-protein interactions, which may lead to inaccurate interactions between the lipids and ESCRT-III proteins (65). We conducted a 200 ns CHARMM36 simulation initialized from one of the backmapped Martini3 tubule structures to see if relaxing into all-atom force field parameters would lead to an adjustment of bilayer thickness. While leaflet peak shapes did change over the course of the 200 ns, there was only a small shift in the leaflet radial positions, suggesting that they are largely fixed over this timescale (Figure S10). The observation that highly curved bicelles from the Martini3 simulations of Mix 1 (Fig. 4D, orange curve) do not reproduce the highly thinned membrane structure we observed in the Martini simulations of the ESCRT-III filament (Fig. 4D, yellow curve) implies that curvature alone is not responsible for this discrepancy.

### Lipid distributions in flat bilayers and curved bicelles

To pinpoint the molecular origin of the thickness discrepancies, we plotted the distributions of the lipid atoms/beads in Mix 1 in flat bilayers and curved bicelles simulated with Martini3 and CHARMM36. Phosphorous atom distributions reflected the thickness differences we observed with multislice simulations (Fig. 5A and 5E). We observed that for Martini3, in both geometries, cholesterol was found deeper in the bilayer, with a significant portion of the hydroxyl oxygens residing at the bilayer center (Fig. 5B and 5F). However, radial distribution profiles in our previous ESCRT filament simulations using Martini2.2 did not show the same propensity for the cholesterol hydroxyl to reside in the bilayer center (33). We therefore analyzed the lipid distributions in our Martini3 ESCRT-III filament simulation and observed that there was more cholesterol localized at the bilayer midplane (Fig. S5B), consistent with the protein-free simulations (Fig. 5).

**Figure 5.**
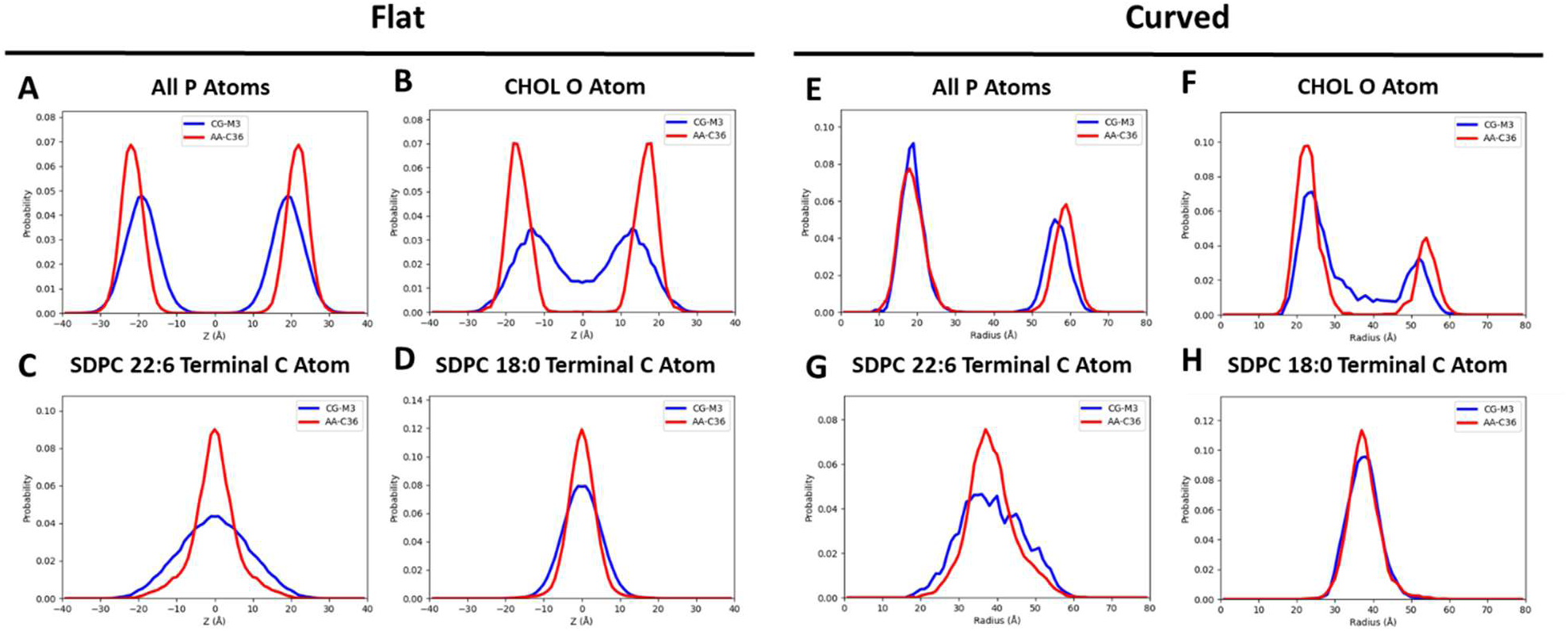
The behavior of cholesterol and polyunsaturated lipids differs strongly between Martini3 and CHARMM36. A-D) Distributions of different atom types reveal differences in the distributions of lipids in Martini- and CHARMM-simulated flat bilayers of Mix 1. E-H) Distributions of different atom types reveal differences in the distributions of lipids in Martini- and CHARMM-simulated curved bicelles of Mix 1.

The reparameterization of Martini cholesterol from version 2 to 3 reduced the free energy barrier for moving the hydroxyl from the hydrated glycerol region to the bilayer center, likely explaining the enrichment of center-residing cholesterol (66, 67). Similarly, the polyunsaturated tail of SDPC was found closer to the bilayer surface in Martini3 simulations compared to CHARMM36 (Fig. 5C and 5G), although both simulations show some degree of backflipping in agreement with previous work (33, 68–73). This behavior was not observed for the saturated tail of SDPC (Fig. 5D and 5H). For cholesterol in hybrid saturated/polyunsaturated PC lipids like SDPC, neutron scattering experiments show cholesterol primarily adopts the canonical upright configuration with little in the bilayer center, which is most consistent with the AA CHARMM36 simulations (74, 75). Both SDPC backflipping and increased cholesterol penetration, along with the hydroxyl, into the bilayer core would be expected to thin the bilayer due to increased phospholipid tail disorder (76, 77). In the Martini3 simulations, removing cholesterol (Mix 3) reduced the SDPC tail backflipping (Fig. S11), and substituting POPC for SDPC (Mix 4) reduced cholesterol localization to the center of the bilayer (Fig. S12), suggesting cooperativity between cholesterol and the 22:6 tail to thin the membrane in Martini simulations. Overall, these data indicated that the lipid bilayer in our ESCRT-III filament Martini simulations was too thin due to unphysical behavior of cholesterol and SDPC, potentially with significant contributions from both bilayer curvature and protein-lipid interactions. Radial distributions calculated from the all-atom filament trajectory showed increased outer leaflet backflipping compared to the protein-free bicelle simulations for both SDPC tails even as cholesterol almost completely reoriented out of the bilayer core, suggesting that protein-lipid tail contact is a primary contributor to thinning (Fig. 5, Fig. S13).

After completion of this work, a reparameterization of the Martini3 lipidome was published (52). This new set of lipid parameters focused on improving phase behavior of complex lipid mixtures, improving the coarse-grained representation of tail length, and expanding the number of parameterized lipids. We repeated the flat bilayer and curved bicelle simulations with the new set of lipid parameters and created simulated cryoEM 2D averages as described for the original Martini3 simulations. Notably, for the lipid mixtures, the new Martini3 parameters (referred to as Martini3.2 here) produced bilayers and bicelles with thicknesses that were more similar to the CHARMM36 simulations (Figs. 6A-C and S11-14). However, for pure POPC, the Martini3.2 simulations produced bilayers and bicelles that were thicker than Martini3, CHARMM36, and the experimental data (Figs. 6D, 3F, and S15). This discrepancy is likely due to the lowering of the average area per lipid that was necessary to improve thermodynamic phase behavior in Martini3.2 and illustrates the compromises that are made when parameterizing coarse-grained force fields. Finally, this example demonstrates the utility of using the analysis pipeline described in this manuscript to evaluate the accuracy of force fields by directly comparing MD simulations to experimental data.

**Figure 6.**
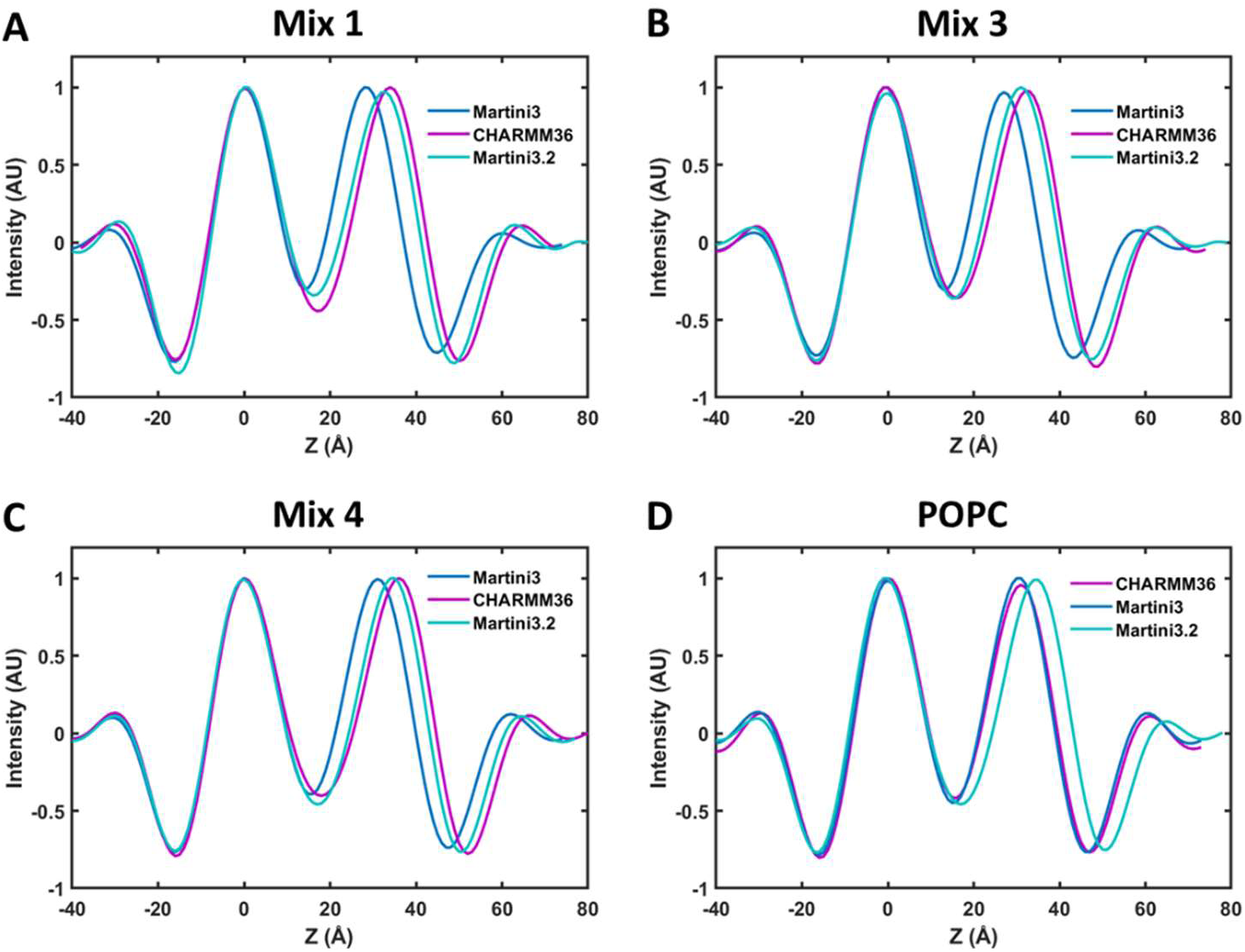
Martini3.2 reparameterized lipids behave more like CHARMM36 lipids for lipid mixtures but not pure POPC. A-D) Comparison of MD simulated bilayer profiles for Mix 1 (A), Mix 3 (B), Mix 4 (C), and POPC (D) shows how force field parameters and lipid mapping schemes affect bilayer thickness.

## Discussion

To test the accuracy of the MD predictions, we converted MD trajectories into simulated cryoEM projections and reconstructed 3D volumes for direct comparison with experimental projections and 3D volumes. With this pipeline, we showed that we can reproduce the low-resolution features of dynamic biological assemblies like lipid bilayers and reproduce the variations in contrast unique to phase contrast TEM imaging. We chose the multislice method to simulate cryoEM projection images due to the accurate, physics-based approach and the ease of including microscope aberrations and the contrast transfer function. However, we note that it does neglect bonded potentials and polarization, which are typically small (78). For computational efficiency, we also excluded the explicit solvent and protons, but these atoms can be included if desired. Additionally, we ignore the physical freezing process, where small changes in molecular conformation are possible (79). Using our pipeline, we compared Martini coarse-grained simulations of a membrane-bound ESCRT-III filament to experimental data. Our synthetic reconstructions pinpointed the limitations of the Martini force field in simulating certain lipids and highly curved protein-bound membrane tubes. Surprisingly, when Martini MD simulations of a membrane-bound ESCRT-III filament were converted to a simulated cryoEM 3D reconstruction, the lipid bilayer was ∼9 Å too thin (Fig. 2E). We do not know whether this discrepancy was due to the setup of the simulation of the CG Martini force field used, but our multislice pipeline identified the inconsistency with experimental data so we could investigate it further. Due to the size of the system, it remains unknown whether the CHARMM36 force field would produce a trajectory that agrees with experimental data.

Heberle and coworkers compared bilayer thicknesses from cryoEM images of SUVs with values obtained from fitting SAXS data and found that cryoEM-derived values were consistently several Å thinner than those obtained by SAXS (29). While these authors did not conclusively determine the cause of this discrepancy, they attributed it to the differences in scattering electrons and X-rays. Here, we showed that the discrepancy is mostly due to anisotropic lipid headgroup peak broadening that makes the membrane appear thinner in cryoEM projection images. In the former case, low resolution cryoEM images result in anisotropic broadening of the leaflet peaks, while the electron density peaks from the higher resolution SAXS data give rise to a greater peak-to-peak difference (Fig. 3D). When both effects are properly accounted for, the multislice workflow accurately predicts the bilayer thicknesses of vesicles of various lipid compositions when compared to SAXS (Fig. 3F). Additionally, neither of these effects explain why the lipid bilayer in the simulated CHMP1B/IST1 reconstruction is thinner than that observed experimentally, suggesting that the difference is due to the CG-MD force field.

To probe this discrepancy, we compared atomistic and CG-MD simulations of lipid bilayers in different geometries with experimental cryoEM data. While Martini and CHARMM simulated membranes sometimes agreed well, we showed that in Martini3 simulations cholesterol tends to localize too deep within the bilayer, while polyunsaturated phospholipid tails are too shallowly inserted into the bilayer (Fig. 5). Further, mixed Martini3 bilayers appear to bend more readily into uniformly curved tubes, while the mixed CHARMM36 simulations kinked (Fig. 4B). For POPC, Martini appeared to perform well in both flat bilayers and curved bilayers. However, the addition of cholesterol to Martini, especially Martini3, simulations caused bilayer thinning compared to AA-MD simulations. This thinning problem was compounded when highly unsaturated lipid tails were added. In these mixtures, the membrane mechanics likely changed as well, as curved bicelles adopted different shapes in Martini3 versus CHARMM36. However, the thinning was even greater in the ESCRT-III copolymer system, and we believe that this could be due to lipid-protein interactions in the coarse-grained systems. The simulated ESCRT-III bound tubule retains an overly thinned outer leaflet with backflipping of both unsaturated and saturated lipid tails even after 200 nanoseconds of relaxation following conversion to all-atom parameters. Cholesterol, however, reorients out of the bilayer center at this timescale. This suggests that overly strong interactions between lipid tails and membrane-inserted protein during the coarse-grained setup drive the excess thinning by decreasing packing in the outer leaflet. Misalignment of coarse-grained cholesterol may also contribute, though that effect appears less robust. It is not clear that further relaxation in all-atom would correct the thinning without adding more lipids and re-equilibrating leaflets.

We expect that the workflow described here will be ideal for directly comparing MD simulations with experimental cryoEM reconstructions to develop a deeper understanding of the physical properties of membranes (especially under high curvature), identify when experiment and simulation fail to match, and facilitate the development of more faithful force fields over a wide range of conditions. Indeed, Martini forcefield optimization remains an active area of research, and we were able to track how changes to the force field impacted our MD simulations as updates were released (Fig. 6) (52, 65, 80). Furthermore, given the exact nature of the forward model for image formation in the TEM, researchers will be able to extract more precise connections between molecular motion and biological mechanisms by comparing cryoEM with MD simulations in the imaging plane.

## Data and Code Availability

MD simulation snapshots and code will be made available on Zenodo.

## Acknowledgements

M.G. and J.L. were supported by NSF grant (MCB 2217662 to M.G.). A.M., A.N., A.F., and F.M. were supported by Altos Labs. A.M., A.F., and F.M. were supported by the NIH (P50 GM082545 to A.F.), and A.F. acknowledges support from the Chan Zuckerberg Biohub. F.M. was supported by a fellowship from the Jane Coffin Childs Memorial Fund for Medical Research. Structural biology applications used in this project were compiled and configured by SBGrid. CryoEM data was collected at the Stanford Cryo-Electron Microscopy Center. We thank Brandon Read and Elizabeth Montabana for assistance with data collection. Work at the Molecular Foundry was supported by the Office of Science, Office of Basic Energy Sciences, of the U.S. Department of Energy under Contract No. DE-AC02-05CH11231. C.O. acknowledges support from the U.S. Department of Energy Early Career Research Award program. Simulations were carried out on the UCSF Wynton Cluster, made possible through grant nos. NIH 1S10OD021596 and 1S10OD020054-011. We thank Dr. Niek van Hilten and Prof. Bob Glaeser for valuable discussion.

## Author Contributions

F.M. and A.F. conceived of and planned the project. A.M., J.L., and A.N. carried out simulations. C.O. wrote simulation code. M.G., A.F., and F.R. supervised research. A.F. and M.G. obtained funding. All authors wrote and edited the manuscript.

