## Supplemental Information for "Comparing Multislice Projections of MD Simulations with CryoEM Exposes Structural Prediction Errors"

**Table of Contents**

1. Simulation methodology
2. Effects of explicit solvent and ions and inclusion of hydrogen atoms on multislice image simulation
3. Simulated projection images, 2D class, averages, and power spectra
4. Sampling a single MD trajectory versus many trajectories
5. Fitting MD snapshots into experimental cryoEM density
6. Martini3 compared to Martini2.2 for ESCRT-III filament simulations
7. MD simulations of protein-free lipid bilayers
8. References

1. **Simulation methodology**

The lipid mixtures used in the simulations are shown in Table S1.

| Mixture Number | Components | Ratio |
|---|---|---|
| 1 | SDPC:POPS:CHOL:PIP2 | 58:18:18:6 |
| 2 | SDPC:POPS:CHOL:PIP2 | 26:32:22:20 |
| 3 | SDPC:POPS:PIP2 | 71:21:8 |
| 4 | POPC:POPS:CHOL:PIP2 | 58:18:18:6 |

**Table S1. Lipid mixtures used in MD simulations.** Ratios shown are molar ratios. SDPC: 1-stearoyl-2-docosahexaenoyl-sn-glycero-3-phosphocholine. POPS: 1-palmitoyl-2-oleoyl-sn-glycero-3-phospho-L-serine. POPC: 1-palmitoyl-2-oleoyl-glycero-3-phosphocholine. CHOL: cholesterol. PIP2: 1,2-dioleoyl-sn-glycero-3-phospho-(1'-myo-inositol-3',5'-bisphosphate).

2. **Effects of explicit solvent and ions and inclusion of hydrogen atoms on multislice image simulation**

We evaluated whether including explicit solvent and solutes in the atomic coordinates used for multislice simulations altered the density of the lipid bilayer, e.g. from ions associated with charged lipid headgroups. As shown in Fig. S1A, there was no significant difference from the inclusion of solvent and solutes. As the inclusion of solvent greatly slows multislice simulations, particularly for such large simulation boxes, we therefore opted to exclude solvent and solutes and to use a Poisson noise model for solvent instead. Similarly, we evaluated whether including non-solvent hydrogen atoms had a significant effect on the multislice projections. Including hydrogens nearly triples the number of atoms, slowing multislice simulations, so we performed this experiment on flat POPC bilayers. As shown in Fig. S1B, inclusion of hydrogens resulted in a small (~1 Å) decrease in apparent bilayer thickness, potentially due to increased electron scatter from the hydrophobic core of the bilayer.

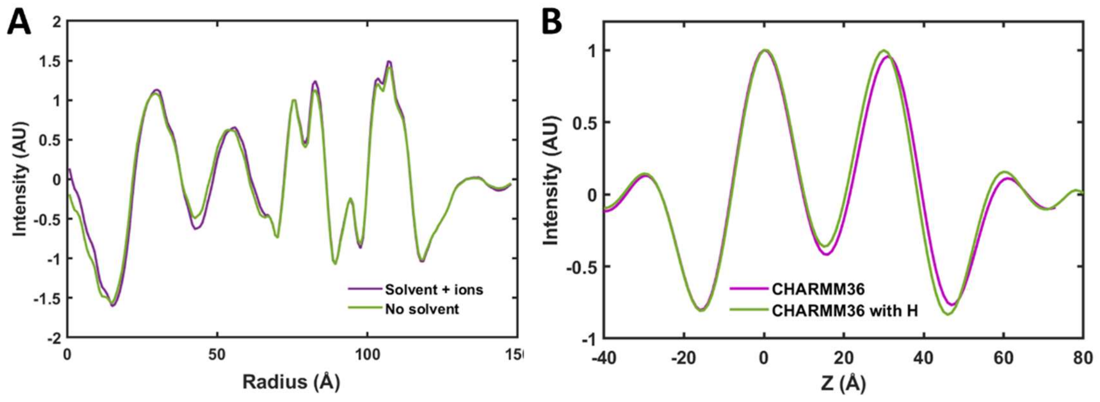

**Figure S1. Comparison of radial potential profiles for multislice simulated cryoEM reconstructions with and without explicit solvent and solutes and with and without hydrogen atoms.** A) Simulated volumes were created from back-mapped Martini2.2 MD simulations as described in the Main Text, except that in one case solvent and solute atoms were not removed. Lipid Mix 1 was used for the membrane. B) Hydrogens were either included or not included from the same CHARMM36 MD simulation snapshots during the multislice projection generation.

### 3. Simulated projection images, 2D class, averages, and power spectra

CryoEM projection images of membrane-bound CHMP1B/IST1 copolymer filaments were randomly selected from the particle stack used to reconstruct the experimental volume (accessed via EMPIAR-11277) (1). Similarly, multislice simulated TEM images of MD simulation snapshots were randomly selected for comparison. 2D averages and power spectra were created in RELION3.1.4.

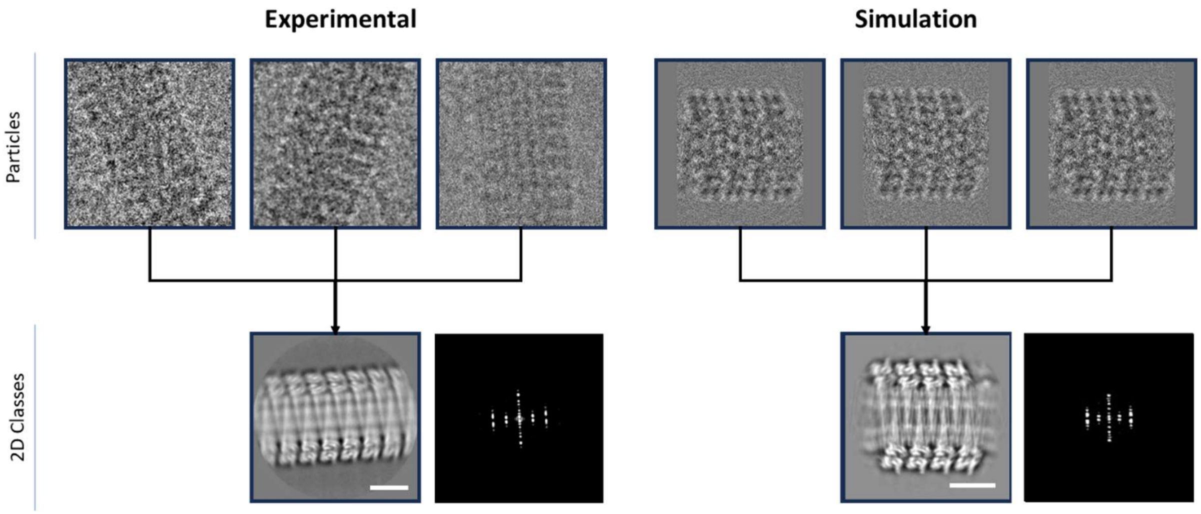


**Figure S2. Multislice simulations produce accurate 2D projection images, 2D averages, and power spectra.** Experimental cryoEM images are shown in 400 Å boxes, while the simulated projection images are shown in 370 Å boxes. Scale bars represent 10 nm.

### 4. Sampling a single MD trajectory versus many trajectories

We wanted to test the ergodicity of our simulations by comparing a single, long trajectory with multiple shorter, independent trajectories. In general, the shorter independent simulations are more efficient and would be preferred as a source of snapshots for the multislice simulation workflow. To improve the computational efficiency of the long MD simulation, we only simulated two turns of the helix, while we simulated four turns in the shorter simulations. Fig. S3 shows that slices of simulated cryoEM volumes generated with each approach are very similar. In both cases, the lipids have decorrelated to form smooth leaflets, indicating that both approaches generate sufficient samples to qualitatively reproduce the membrane structure from the experimental data.

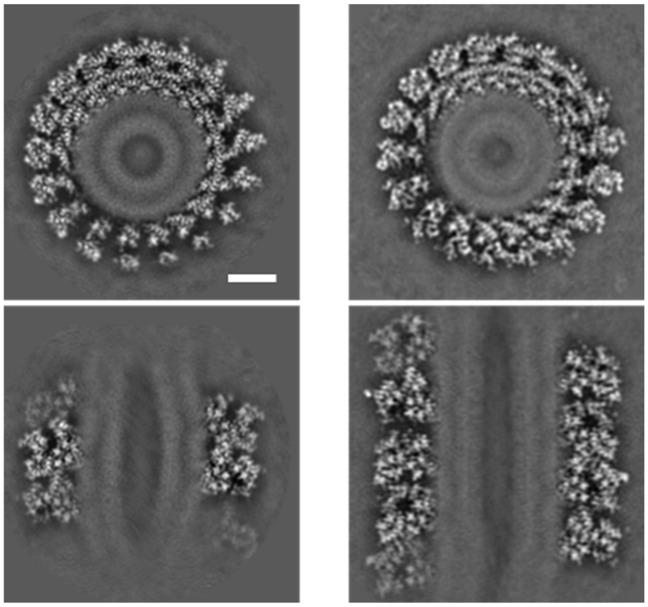

**Figure S3. Comparison of MD sampling strategies.** Horizontal (top) and vertical (bottom) slices from multislice simulated cryoEM volumes generated using a single, two-turn helical filament trajectory sampled at 10 time points (left) compared to four-turn helical filament sampled from 15 independent simulations (right). Scale bar represents 5 nm.

### 5. Fitting MD snapshots into experimental cryoEM density

A representative all-atom MD snapshot back-mapped from a Martini2.2 simulation was fit into the experimental cryoEM density (accessed from EMD-28700 and low pass filtered to 6 Å) using the Fit in Map function in Chimera (Fig. S4). The protein atoms fit into the density well. However, it is unclear how well the lipids fit into the two bands of density for the inner and outer leaflets of the bilayer.

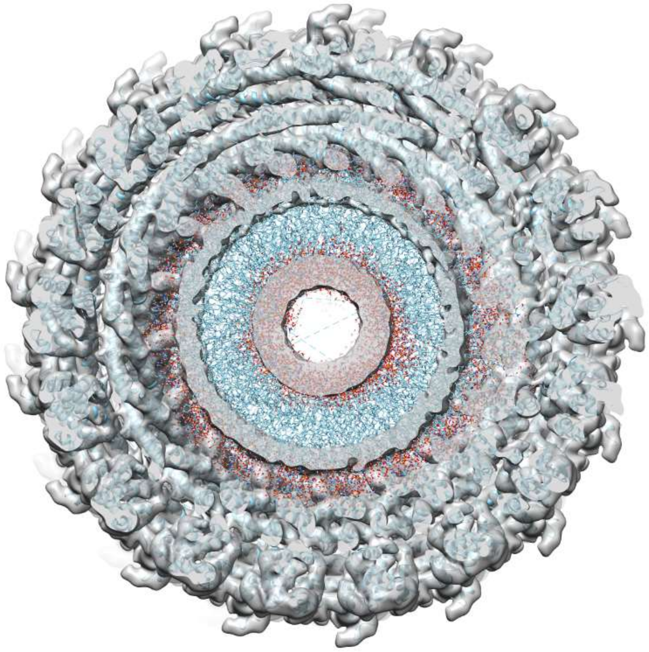

**Figure S4. Fitting of an MD snapshot into experimental cryoEM density for the membrane-bound CHMP1B/IST1 copolymer filament.** The protein is displayed as a ribbon diagram in cyan, and lipid atoms are displayed with a stick model, colored according to atom type. Carbon is cyan, oxygen is red, nitrogen is blue, and phosphorous is orange.

### 6. Martini3 compared to Martini2.2 for ESCRT-III filament simulations

Martini3 was released after performing many of the MD simulations with Martini2.2. We therefore wanted to evaluate whether the updated parameters in Martini3 changed the bilayer structure in our simulations. We directly compared radial potential profiles from simulated volumes generated from Martini2.2 and Martini3 CG-MD simulations. As shown in Fig. S5A, Martini3 matched Martini2.2 closely, with Martini3 generating a slightly thinner bilayer. Considering our result that Martini3 places cholesterol too deeply within protein-free bilayers, we also examined the distribution of the cholesterol OH bead. Indeed, this bead was more likely to be found near the center of the bilayer than in Martini2.2 simulations (Fig. S5B).

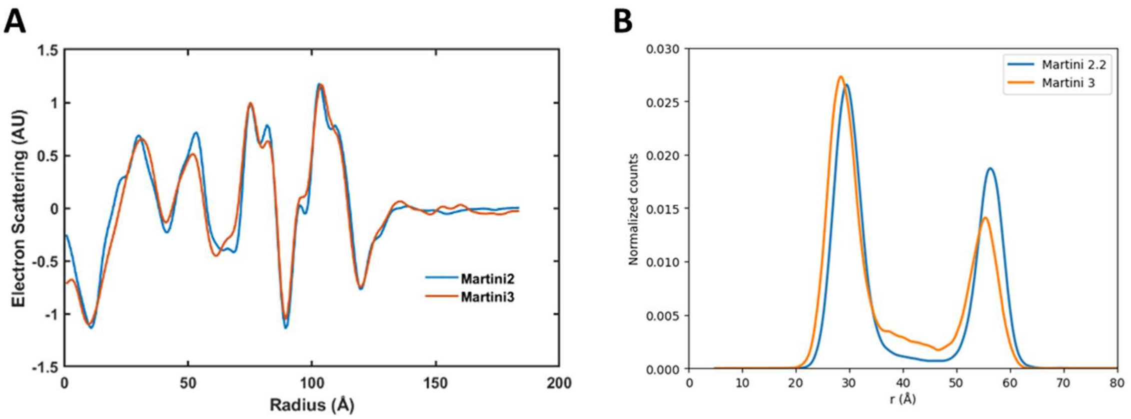


**Figure S5. Comparison between Martini2.2 and Martini3 CG-MD simulations of the membrane-bound CHMP1B/IST1 copolymer filament.** A) Radial potential profiles of multislice simulated cryoEM volumes generated from MD simulations utilizing the two forcefields. B) Radial distributions of cholesterol OH beads during MD simulations performed with the two forcefields.

### 7. MD simulations of protein-free lipid bilayers

As described in the Main Text, we empirically determined the low-pass filter necessary to match the resolution of the experimental 2D average for POPC. The full range of low-pass filtered simulated 2D averages is shown in Fig. S6. 20-30 Å low-pass filtering provided the best correspondence.

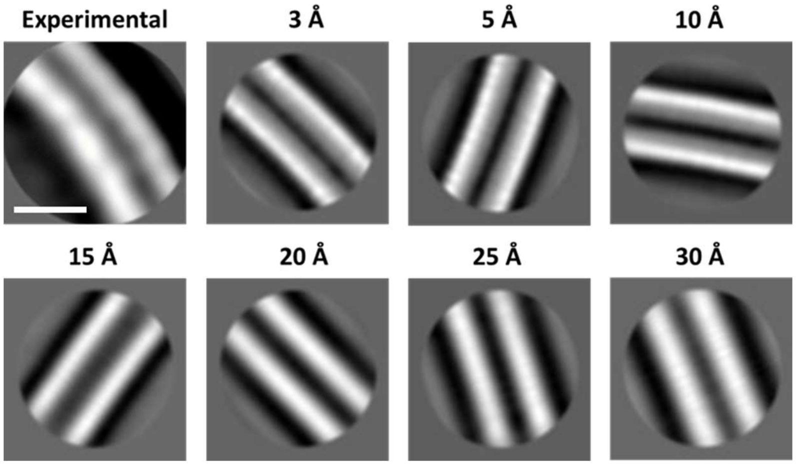


**Figure S6. Experimental and multislice simulated 2D averages of POPC bilayers at various levels of low-pass filtering.** Scale bar represents 5 nm.

| POPC Bilayer | Thickness (Å) |
|---|---|
| Simulated SAXS | 38 |
| Experimental SAXS | 36.3 |
| Experimental TEM | 29.4 |
| Simulated 3 A TEM | 35 |
| Simulated 30 A TEM | 32 |

**Table S2. Measured peak-to-peak thicknesses from experimental and simulation for POPC bilayers.**

The full set of 2D averages and additional Coulombic potential profiles not shown in the Main Text are shown in Fig. S7. Refer to Table S1 for lipid compositions. Additionally, the measured peak-to-peak bilayer thicknesses calculated from 2D averages and SAXS data of POPC are shown in Table S2.

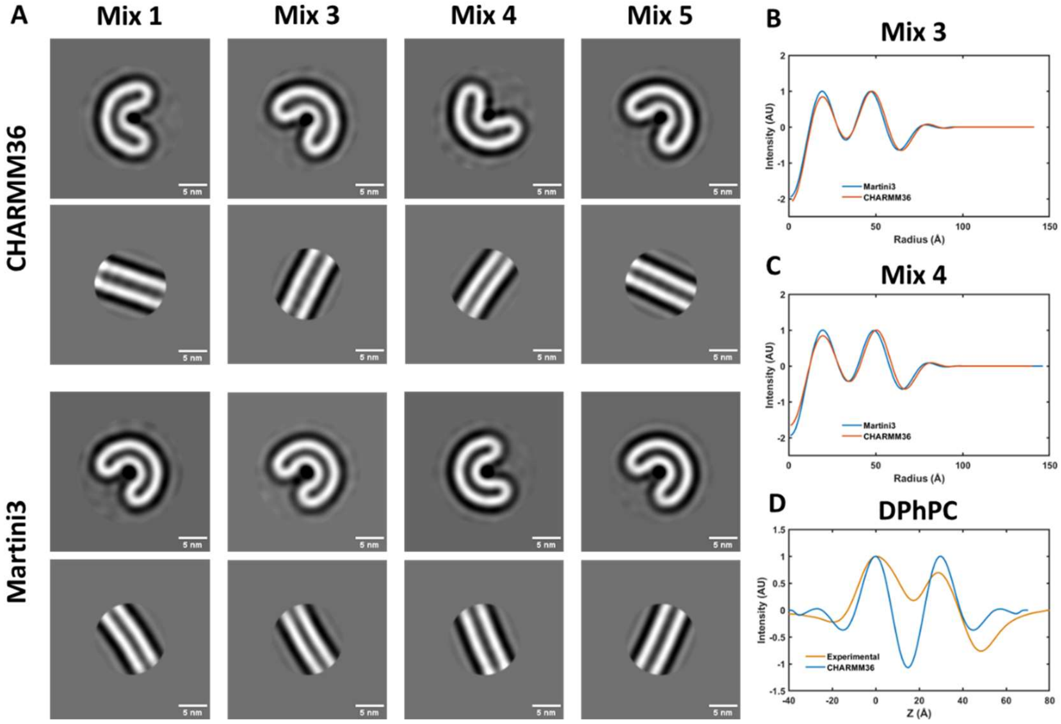


**Figure S7. Comparison of flat bilayers and curved bicelles of different lipid compositions.** A) Simulated 2D class averages of four lipid mixtures for flat bilayers and curved bicelles simulated with Martini3 and CHARMM36. Scale bar = 5 nm. B) radial and linear potential profiles for Martini3 and CHARMM36 simulated bicelles and bilayers.

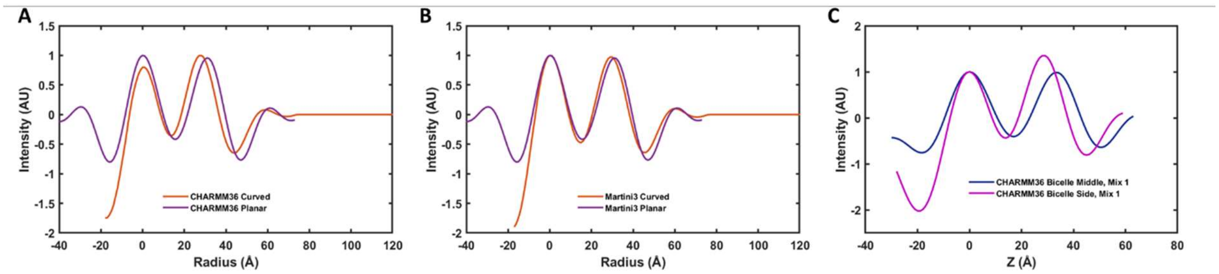


**Figure S8. Curved bicelles are thinner than flat bilayers.** A) Comparison of profiles of flat and planar POPC bilayers and bicelles simulated with CHARMM36. B) Comparison of profiles of flat and planar POPC bilayers and bicelles simulated with Martini3. C) Comparison of the membrane thickness in the middle and at the sides of the Mix 1 bicelle.

We also computed difference maps between CHARMM36 and Martini3 simulations of curved bicelles and flat bilayers. Consistent with our measurement from multislice simulations, these difference maps show that CHARMM36 results in a thicker membrane than Martini3 for lipid mixtures 1, 3, and 4, but not pure POPC. The differences at the edges of the bicelles also demonstrate the differing geometries predicted by these forcefields. Note that the difference maps were generated directly from electron density in the MD simulation snapshots, not using multislice methods.

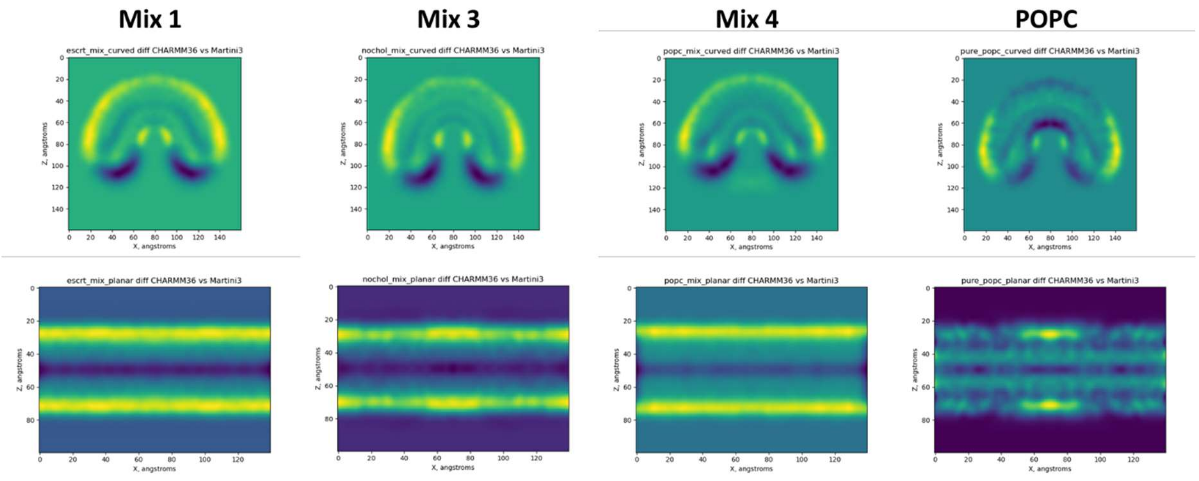


**Figure S9. Difference maps calculated as the electron density from CHARMM36 snapshots minus that from Martini3 snapshots for curved bicelles (top) and flat bilayers (bottom).**

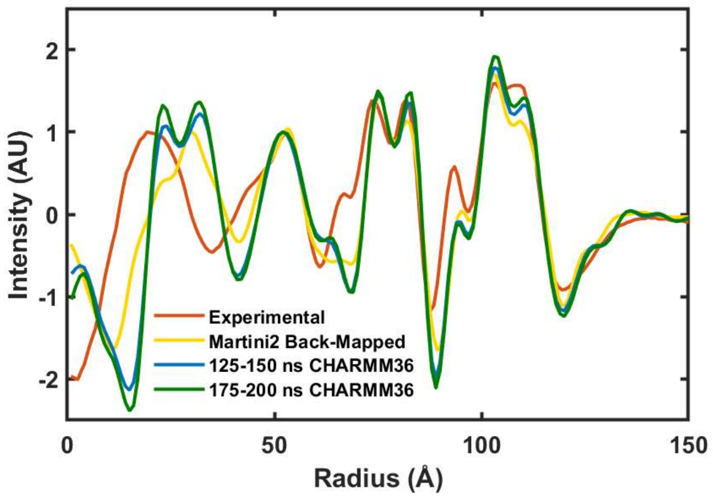


**Figure S10. Radial potential profiles of experimental and simulated protein-coated lipid nanotubes.** Comparison of the experimental EM profile with simulation set 3 Martini2 back-mapped profile and with profiles extracted from extended all-atom tubule simulation initialized from a backmapped Set 4 Martini3 structure.

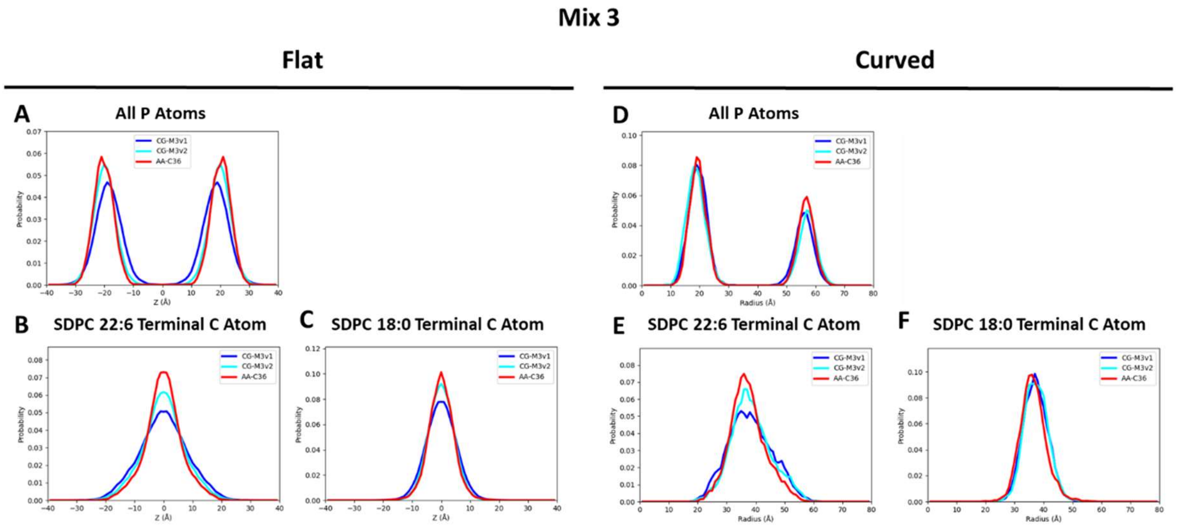


**Figure S11. Distributions of different atom and bead types in CHARMM36 and Martini3 simulations of flat bilayers and curved bicelles composed of Mix 3, as in Fig. 5.** Distributions of different atom types reveal differences in the distributions of lipids in Martini- and CHARMM-simulated flat bilayers (A-C) and curved bicelles (D-F) of Mix 3.

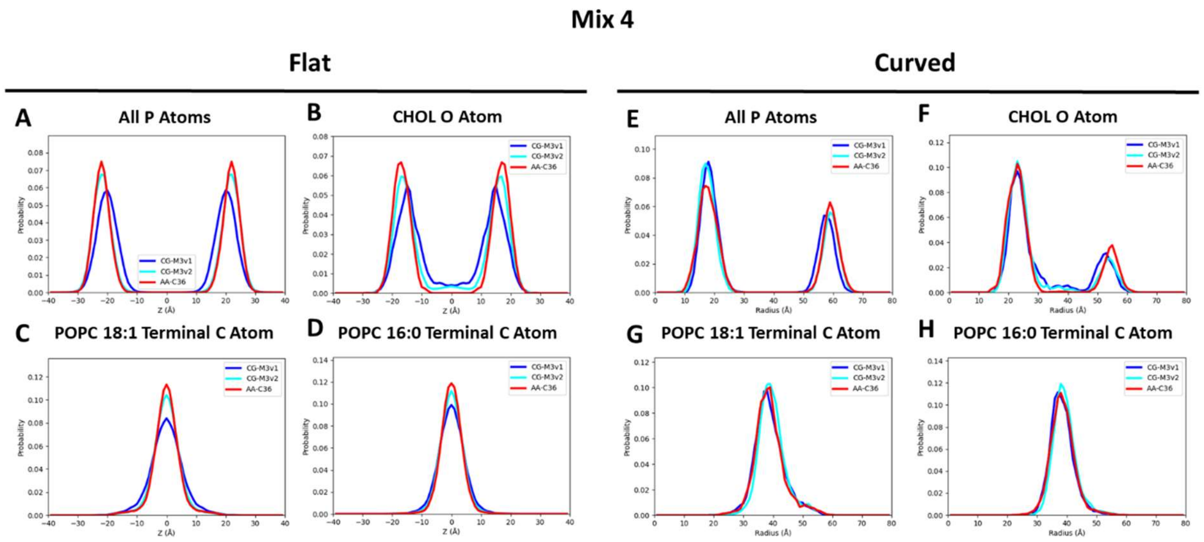


**Figure S12. Distributions of different atom and bead types in CHARMM36 and Martini3 simulations of flat bilayers and curved bicelles composed of Mix 4, as in Fig. 5.** Distributions of different atom types reveal differences in the distributions of lipids in Martini- and CHARMM-simulated flat bilayers (A-D) and curved bicelles (E-H) of Mix 4.

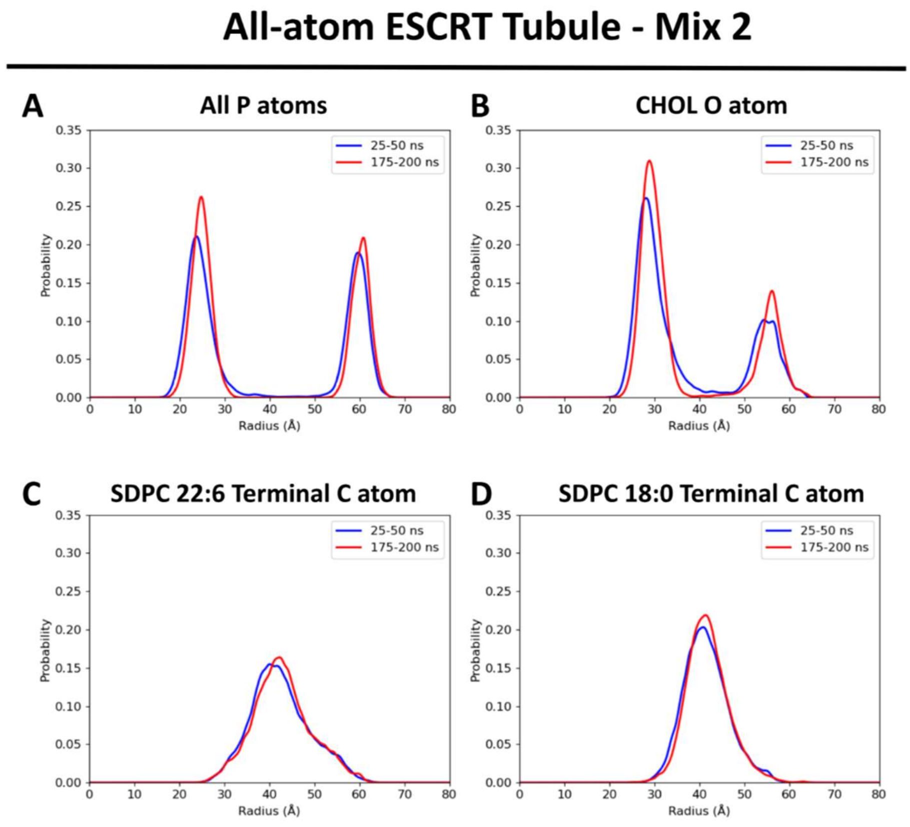

**Figure S13. Distributions of different atoms types in CHARMM36 from all-atom ESCRT Tubule trajectory, as in Fig. 5.** Radial distributions calculated from the 25-50 ns segment and 175-200 ns segment of the all-atom ESCRT tubule trajectory initialized from a simulation Set 4 structure, after back-mapping from Martini3. Differences between the two sets reflect the extent to which the tubule can relax to the new force field, primarily in sharpening of phosphate/oxygen peaks and displacement of cholesterol from the bilayer center.

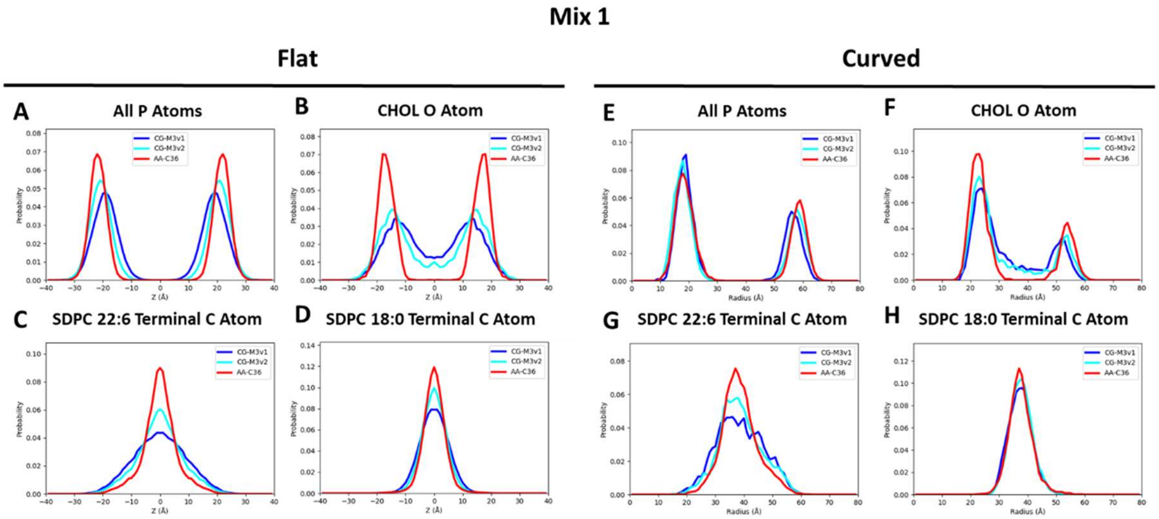


**Figure S14. Distributions of different atom and bead types in CHARMM36, Martini3, and Martini3.2 simulations of flat bilayers and curved bicelles composed of Mix 1, as in Fig. 5.** Distributions of different atom types reveal differences in the distributions of lipids in Martini- and CHARMM-simulated flat bilayers (A-D) and curved bicelles (E-H) of Mix 1.

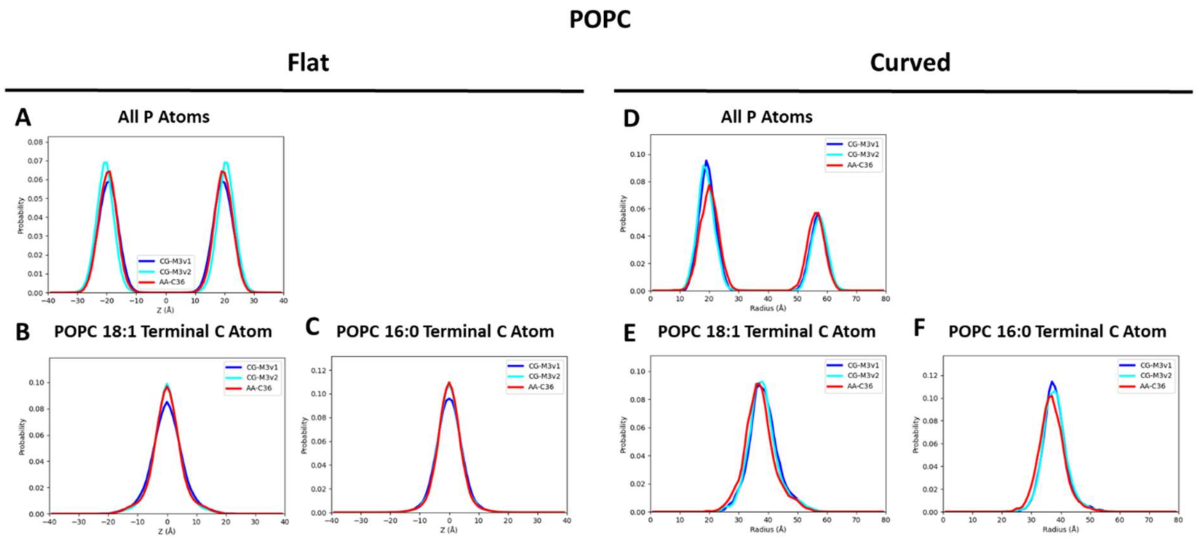


**Figure S15. Distributions of different atom and bead types in CHARMM36, Martini3, and Martini3.2 simulations of flat bilayers and curved bicelles composed of POPC, as in Fig. 5.** Distributions of different atom types reveal differences in the distributions of lipids in Martini- and CHARMM-simulated flat bilayers (A-D) and curved bicelles (E-H) of POPC.
